# Evolution and diversification of the momilactone biosynthetic gene cluster in the genus *Oryza*

**DOI:** 10.1101/2024.01.11.572147

**Authors:** Santiago Priego-Cubero, Tomonobu Toyomasu, Michael Gigl, Youming Liu, Yuto Hasegawa, Hideaki Nojiri, Corinna Dawid, Kazunori Okada, Claude Becker

## Abstract

Plants are master chemists and collectively are able to produce hundreds of thousands of different organic compounds. The genes underlying the biosynthesis of many specialized metabolites are organized in biosynthetic gene clusters (BGCs), which is hypothesized to ensure their faithful co-inheritance and to facilitate their coordinated expression. In rice, momilactones are diterpenoids that act in plant defence and various organismic interactions. Many of the genes essential for momilactone biosynthesis are grouped in a BGC. Here, we apply comparative genomics of diploid and allotetraploid *Oryza* species to reconstruct the species-specific architecture, evolutionary trajectory, and sub-functionalisation of the momilactone biosynthetic gene cluster (MBGC) in the *Oryza* genus. Our data show that the evolution of the MBGC is marked by lineage-specific rearrangements and gene copy number variation, as well as by occasional cluster loss. We identified a distinct cluster architecture in *O. coarctata*, which represents the first instance of an alternative architecture of the MBGC in *Oryza* and strengthens the idea of a common origin of the cluster in *Oryza* and the distantly related genus *Echinochloa*. Our research illustrates the evolutionary and functional dynamics of a biosynthetic gene cluster within a plant genus.

## INTRODUCTION

### Biosynthetic gene clusters and their evolution

With more and more plant reference genome assemblies becoming available, biosynthetic gene clusters (BGCs), i.e., the co-localization of often phylogenetically unrelated genes that participate in the same biosynthetic cascade of specialized metabolites, have emerged as a common feature of genomic organization in plants (Polturak et al., 2022b). BGCs are postulated to confer evolutionary advantages because they facilitate coordinated gene expression, enable the reliable coinheritance of genes involved in the same metabolic pathway (thereby preventing the accumulation of toxic intermediates), or facilitate the formation of metabolons (Nützmann et al., 2016). However, the mechanisms by which such non-orthologous genes become localized in the same genomic region and act in the same biosynthetic pathway are still poorly understood. Currently, the most common model proposes that they have formed through a series of events that is driven by both positive and negative selection pressure, starting with gene duplication, followed by neo-functionalization, and ultimately relocation. In some cases, this process appears to have been mediated by transposable elements (Polturak et al., 2022b; Smit and Lichman, 2022).

### Biological functions of rice phytoalexins, labdane-related diterpenoids and momilactones

Phytoalexins are low-molecular-mass specialized plant metabolites that are often produced under biotic and abiotic stress conditions (Ahuja et al., 2012). In rice (*Oryza sativa*), the major phytoalexins are a group of labdane-related diterpenoids (reviewed in Toyomasu et al., 2020), which derive from the cyclization of geranylgeranyl diphosphate (GGPP) into *ent*, *syn*, or normal stereoisomers of copalyl diphosphate (CDP) by the class II diterpene synthases Copalyl Diphosphate Synthases (CPSs). The biosynthesis of these metabolites has evolved from that of gibberellins (GAs), *ent* labdane-related diterpenoids themselves, through duplication and neofunctionalization of core biosynthetic enzymes (Zi et al., 2014). Several *ent* and *syn* (but not normal) rice labdane-related diterpenoids have been identified, including momilactones A and B, phytocassanes A to F, and oryzalexins (A to F, and S) (Toyomasu et al., 2020; Zi et al., 2014). Notably, momilactone A and, more prominently, momilactone B have a strong allelopathic activity, i.e., they inhibit the germination and growth of nearby plants upon being released by the rice plants into the soil (Kato et al., 1973; Kato-Noguchi et al., 2010; Serra Serra et al. 2021). Both compounds accumulate in rice husks but are also exuded from the roots (Kato-Noguchi et al., 2010; Kato-Noguchi and Ino, 2003).

### Biosynthesis of momilactones and clustering of momilactone genes

Momilactone biosynthesis starts with the cyclization of GGPP into *syn*-copalyl diphosphate (*syn*-CDP), catalysed by the Copalyl Diphosphate Synthase 4 (CPS4) (Otomo et al., 2004b; Xu et al., 2004). *syn*-CDP is further cyclized into *syn*-pimara-7,15-diene by *ent*-kaurene synthase-like 4 (KSL4), a class I diterpene synthase (Otomo et al., 2004a; Wilderman et al., 2004). Because *syn*-CDP is also a substrate for oryzalexin S biosynthesis, the KSL4-mediated cyclization is the first truly dedicated step towards momilactone production (Tamogani et al., 1993). 9βH-pimara-7,15-diene undergoes several oxidation steps, catalysed first by cytochrome P450 (CYP) monooxygenases CYP76M8 and CYP99A3, followed by the short-chain alcohol dehydrogenase MOMILACTONE A SYNTHASE (MAS) and CYP701A8, to yield momilactone A (De La Peña and Sattely, 2021; Kitaoka et al., 2021). CYP76M14 catalyses the final hydroxylation of C20, leading to spontaneous closure of the hemi-acetal ring and forming momilactone B (De La Peña and Sattely, 2021). Among the biosynthetic genes, *CPS4*, *KSL4*, and the paralogs *CYP99A2/3* and *MAS1/2* are co-localized on chromosome 4 in a momilactone biosynthetic gene cluster (MBGC) (Miyamoto et al., 2016; Shimura et al., 2007). However, not all genes required for momilactone biosynthesis are contained in the MBGC: *CYP76M8* is located on chromosome 2 and is part of another BGC that is required for phytocassane and oryzalexin production (Okada, 2011; Kitaoka et al., 2021), while *CYP701A8* and *CYP76M14* are located on chromosomes 6 and 1, respectively (De La Peña and Sattely, 2021).

### Evolution of the momilactone biosynthetic gene cluster in Oryza

The *Oryza* genus (belonging to the *Poaceae* family) consists of 27 known species with 11 different genome types (classified based on cytogenetics and genetic hybridization studies): 6 diploids (n = 12; AA, BB, CC, EE, FF and GG) and 5 allotetraploids (n = 24; BBCC, CCDD, HHJJ, HHKK and KKLL) (Ge et al., 1999; Lu et al., 2009). The MBGC was established in *Oryza* before the domestication of rice (*O. sativa*; AA) (Miyamoto et al., 2016). It is highly conserved among AA and BB genome type *Oryza* species, while only a partial cluster exists in the early-divergent *Oryza* species *O. brachyantha* (FF), which harbours only two clustered *CYP99A2/3* paralogs (Miyamoto et al., 2016). Because the MBGC is incomplete in *O. brachyantha*, it likely evolved prior to the divergence of the BB lineage in *Oryza* (Miyamoto et al., 2016). Due to the limited availability of genome assemblies from species positioned between *O. brachyantha* and *O. punctata* in the phylogenetic tree, it has remained unclear whether the MBGC was restricted to species of the AA and BB lineages. Alternatively, the MBGC could have been lost specifically in the *O. brachyantha* lineage and might be present in other *Oryza* lineages. Lastly, the presence and configuration of the MBGC have not yet been studied in any of the allotetraploid *Oryza* species, and it therefore remains unknown if the cluster is present in both their sub-genomes.

### Momilactone biosynthetic gene cluster in other species

Studies of the MBGC have not been limited to the *Oryza* genus; some have expanded into the wider *Poaceae* family. *Poaceae* encompass twelve subfamilies, of which nine belong to two core clades: BOP and PACMAD. The BOP clade comprises Bambusoideae, Oryzoideae (including *Oryza*), and Pooideae (including the *Triticeae* tribe); the PACMAD clade consists of Panicoideae, Aristidoideae, Chloridoideae, Micrairoideae, Arundinoideae, and Danthonioideae (Soreng et al., 2017). The MBGC is not restricted to the *Oryza* genus and has been identified through phylogenomic methods in species belonging to the PACMAD clade, specifically in the Panicoideae and Chloridoideae subfamilies (Wu et al., 2022a). However, it remains uncertain whether these species actually produce momilactones or other types of labdane-related diterpenoids. *Echinochloa crus-galli* (Panicoideae), a prevalent weed associated with rice cultivation, is the only species confirmed to produce momilactone A (Kraehmer et al., 2016; Wu et al., 2022a). Interestingly, *E. crus-galli* exhibits a distinct MBGC architecture, including an additional cytochrome P450 (*EcCYP76L11*) (Guo et al., 2017; Kitaoka et al., 2021). EcCYP76L11, similarly to CYP76M8 in *Oryza sativa*, can catalyse the conversion of *syn*-pimaradiene into 6β-hydroxy-*syn*-pimaradiene (Kitaoka et al., 2021). Despite the shared origin of the MBGC in *Oryza* and *Echinochloa*, the *CYP76L11* gene has not been identified in the MBGC or the genome of any *Oryza* species (Wu et al., 2022a).

Here, we studied the presence, architecture, and evolution of the MBGC in cultivated rice and wild relatives from the *Oryza* genus by mining recently published genome assemblies. Our study shows that the MBGC is not restricted to species of the AA and BB lineages; we identify MBGC-like clusters in *Oryza* CC species and in one of the respective sub-genomes of the tetraploid species *O. alta* as well as in *O. coarctata*, a basal lineage in the *Oryza* phylogeny. We also show that the gene cluster was lost in several intermediate genome types. While momilactone A was detectable in all species harbouring the cluster, our data indicate that momilactone B production might have been a more recent innovation that emerged after the branching off of the KK genome type. In the *O. coarctata* MBGC, we identified an additional CYP that is different from the canonical *O. sativa* CYP99A2/3 and for which a corresponding ortholog could be found only in the MBGC of *E. crus-galli*, altogether suggesting the existence of an early ancestral cluster. In summary, our study shows how a biosynthetic gene cluster diversified within a plant genus while largely maintaining its biosynthetic function.

## RESULTS

### The momilactone biosynthetic gene cluster is prevalent throughout the Oryza genus

Among the 11 different genome types within the genus *Oryza*, the MBGC has been extensively studied in only 3 lineages. The cluster exhibits a high level of conservation among *Oryza* species of the AA and BB genome groups, whereas it is incomplete in the FF species *O. brachyantha* (Figure 1b) (Miyamoto et al., 2016). To better understand the evolution of MBGC in the different *Oryza* lineages, we mined newly published genome assemblies (Supplementary Table 1) for the presence and location of the MBGC (see methods for details). Our analysis included genomes of the diploid species *O. officinalis*, *O. eichingeri*, *O. rhizomatis* (all CC), *O. australiensis* (EE), and *O. granulata* (GG), as well as of the allotetraploid species *O. alta* (CCDD) and *O. coarctata* (KKLL). We also included *Leersia perrieri* as an outgroup to *Oryza* in our analyses. We used OrthoFinder (Emms and Kelly, 2019) to identify orthogroups containing putative *O. sativa* momilactone biosynthetic genes (Figure 1a). Based on phylogenetic relationships between the set of genes in each orthogroup, we considered as homologs those genes that belonged to the same clade as the *O. sativa* momilactone biosynthetic genes (Supplementary Figures 1-4). We further mined the genomic regions upstream and downstream of the identified homologs to confirm their organization in a genomic cluster.

**Figure 1.**
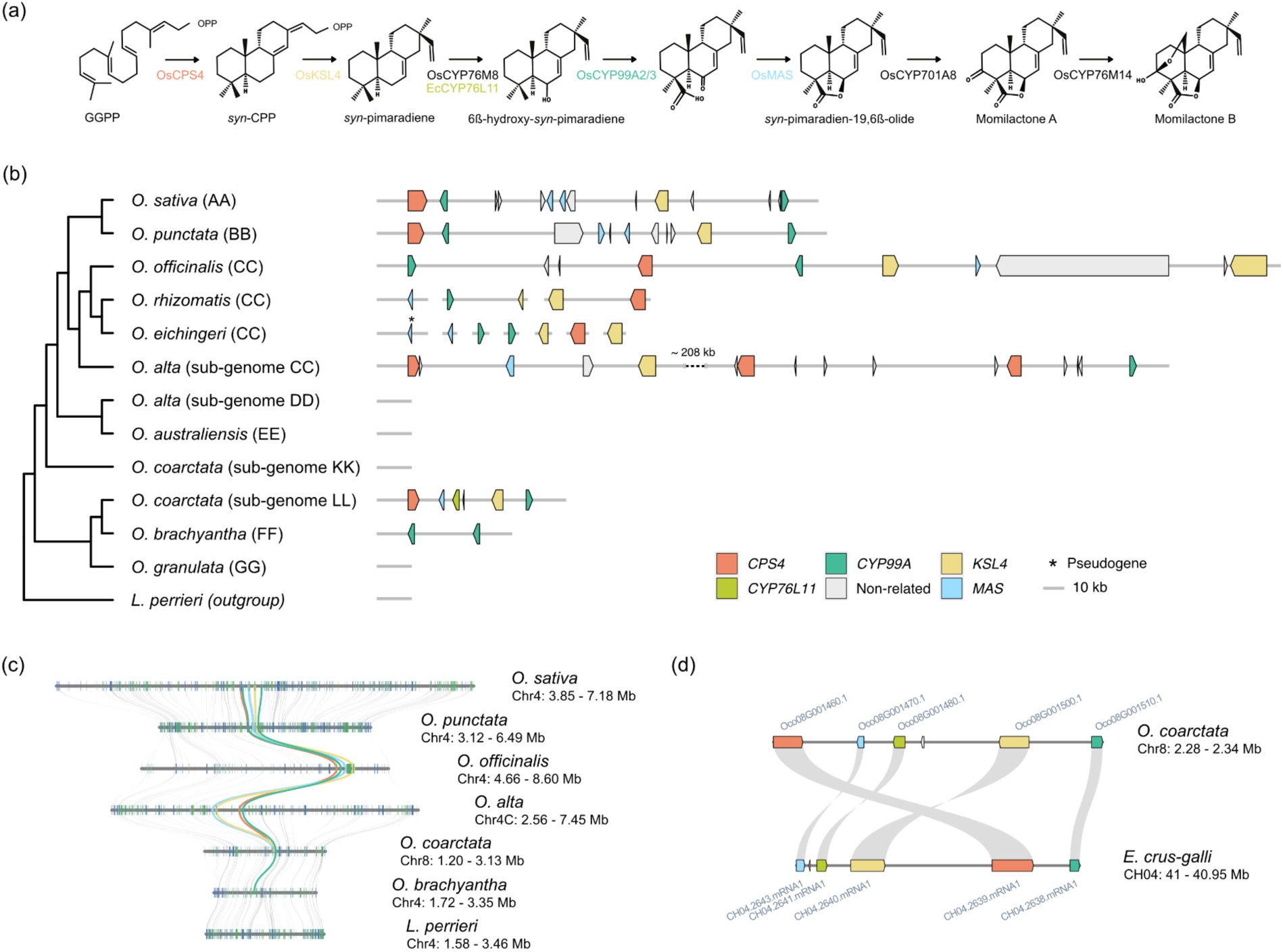
The momilactone biosynthetic gene cluster in the *Oryza* genus. **a)** Simplified representation of the momilactone biosynthetic pathway, adapted from De la Peña et al. (2021) and Kitaoka et al. (2021). **b)** Phylogenetic relationship between the different *Oryza* species and sub-genomes included in this study, the outgroup *L. perrieri*, and their respective MBGCs. The species tree represents the maximum-likelihood tree inferred from a concatenated multiple-sequence alignment of single-copy orthologs using 4,069 orthogroups with a minimum of 100.0% of species having single-copy genes in any orthogroup. Coloured block arrows represent genes. Scaffold and positional information on all genes is provided in Supplementary Table 2. **c)** Microsynteny between the genomic regions containing the MBGC from *O. sativa*, *O. punctata*, *O. officinalis*, *O. alta*, *O. coarctata*, *O. brachyantha*, and *L. perrieri*. **d)** Microsynteny between MBGC in *O. coarctata* and *E. crus-galli*. Note that this microsynteny does not extend to the flanking regions of the cluster (also see Supplementary Figure 6). In (c) and (d), grey lines connect the respective orthologs. “Non-related” refers to genes that do not seem to be functionally related to the main biosynthetic cascade of the cluster.

We detected the MBGC in the CC genome species *O. officinalis*, and the corresponding genes across different scaffolds of two other CC species, *O. eichingeri*, and *O. rhizomatis*. Within the CC lineage, gene copy numbers varied, and for some genes we found signatures of species-dependent duplications. In *O. officinalis* (CC), the cluster (∼385 kb in length) was located on chromosome 4 and contained *CPS4*, *MAS*, two copies of *KSL4*, and two *CYP99A2/3* orthologs (Figure 1b). In *O. eichingeri* and *O. rhizomatis* (both CC), we identified orthologs of *CPS4*, *MAS*, two copies of *KSL4*, and *CYP99A2/3*; *O. eichingeri* carried two copies each for *CYP99A2/3* and *MAS*, with the second copy of the latter appearing pseudogenic (Figure 1b). However, in the scaffold-level assemblies of *O. eichingeri* and *O. rhizomatis*, we found these genes localized on different scaffolds. In *O. rhizomatis*, the *CYP99A2/3* and one of the *KSL4* paralogs co-occurred on Scaffold2280, *CPS4* and the other *KSL4* paralog on Scaffold1491 (Figure 1b). In agreement with the presence of the cluster in all CC species, we identified a ∼520 kb MBGC cluster on chromosome 4 of sub-genome C of the allotetraploid *O. alta* (CCDD). This cluster contained orthologs of *MAS*, *KSL4*, *CYP99A2/3*, and three orthologs of *CPS4* (Figure 1b). In contrast, we did not find any orthologs of momilactone biosynthetic genes in the D sub-genome of *O. alta*. This observation is in line with *O. australiensis* (EE), the closest relative of the *O. alta* D-type sub-genome lacking a MBGC (the ancestral diploid D-type donor is presumably extinct) (Bao and Ge, 2004; Ge et al., 1999). In the allotetraploid *O. coarctata* (KKLL), we identified a MBGC spanning ∼55 kb and containing one ortholog each of *CPS4*, *MAS*, *KSL4*, and *CYP99A2/3*. Based on a species tree obtained from 4,069 multiple-sequence alignments of single-copy orthologs (Figure 1b) and from existing literature (Ge et al., 1999; Guo and Ge, 2005; Shenton et al., 2020), we could tentatively assign the MBGC in *O. coarctata* to chromosome 4 (LG08) of the LL sub-genome (evenly numbered chromosomes in the annotation belong to the LL, unevenly numbered ones to the KK sub-genome). Because this lineage diverged earlier than the MBGC-lacking EE lineage (Figure 1b), we speculate that the cluster most likely became lost in the EE lineage. Lastly, we did not detect the cluster or homologs thereof in *O. granulata* (GG).

To further confirm that we had detected the MBGC and not another diterpene-related BGC, we analysed the synteny of the corresponding regions among the different species. We hypothesized that if the MBGC in the *Oryza* genus had a common origin, one should observe high synteny of the genomic region among all species. We excluded *O. rhizomatis* and *O. eichingeri* from the synteny analysis, because in those genome assemblies the momilactone biosynthesis genes were located on short scaffolds that often carried only very few (∼3-4) genes in total. For the remaining species and sub-genomes, the genomic region containing the MBGC (or lacking it, as in the case of *O. alta* sub-genome D, *O. australiensis*, *O. coarctata* sub-genome K, and *O. granulata*) was syntenic (Figure 1c, Supplementary Figure 5). The synteny further confirmed the presence of the MBGC in the different reference genomes and indicated that this region shares a common evolutionary origin. Taken together, these results suggest that the MBGC is prevalent at a basal position of the *Oryza* phylogeny. The evolution of the cluster in different lineages is characterized by differences in cluster size, lineage-specific rearrangements with an important variation in gene copy number, and lineage-specific loss of the cluster.

### Species-specific deviations in the MBGC architecture in the Oryza genus

We next investigated whether there were any noticeable species-specific deviations in the MBGC architecture. In *O. coarctata*, we identified an extra gene located between *MAS* and *KSL4* and encoding a CYP. Importantly, this enzyme does not belong to the CYP99A subfamily, which is the canonical CYP in the MBGC of the other *Oryza* species (Figure 1b). Instead, the MBGC in *O. coarctata* strikingly resembled that of *E. crus-galli* (Figure 1d), which also contains an additional CYP-encoding gene, *EcCYP76L11*, that does not belong to the CYP99A subfamily either (Guo et al., 2017; Kitaoka et al., 2021). EcCYP76L11 catalyses the same reaction as OsCYP76M8, converting *syn*-pimaradiene to 6β-hydroxy-*syn*-pimaradiene (Kitaoka et al., 2021). In rice, *OsCYP76M8* is located outside of the MBGC, which suggests that *E. crus-galli* possibly contains a more compacted version of the cluster (Kitaoka et al., 2021).

We further investigated the MBGC in *O. coarctata*. An initial microsynteny comparison of the MBGCs in *O. coarctata* and *E. crus-galli* revealed that the additional CYP-encoding gene could indeed be an ortholog of *EcCYP76L11* (Figure 1d). Notably, the microsynteny pattern was restricted to the MBGC itself and not to the flanking regions of the cluster in both species (Supplementary Figure 6). We confirmed that this additional CYP-encoding gene was a reciprocal best hit to EcCYP76L11. To verify the relationship between the CYP-encoding gene in *O. coarctata* MBGC and *EcCYP76L11*, we conducted an amino-acid-sequence-based phylogenetic analysis including all *Oryza* species from our panel and different representative species of the main Poaceae subfamilies (Supplementary Figure 7; see Material and Methods for details). Our analysis placed the *O. coarctata* CYP (Oc08G001480) in a clade with EcCYP76L11 (CH04.2641), the two CYP76L11 paralogs from the MBGC of *E. haploclada* (eh_chr4.2731 and eh_chr4.2733) (Wu et al. 2022) and two CYPs from *P. hallii* (Pahal.1G039100) and *S. italica* (Seita.2G145700), respectively. Interestingly, the *O. coarctata* CYP did not group in a clade with the unique CYP76L member from *O. sativa* (OsCYP76L1) or the CYP76L1 orthologs in other *Oryza* species (including Oc18G006310 from *O. coarctata*) (Figure 2). Consequently, based on the similarity to EcCYP76L11, we named the newly identified *O. coarctata* CYP-encoding gene *OcCYP76L11*. Our analysis suggests that *OcCYP76L11* was present in the ancestral *Oryza* MBGC and was retained in the early-divergent *O. coarctata*, while it became lost in the later-diverging clades of the *Oryza* genus.

**Figure 2.**
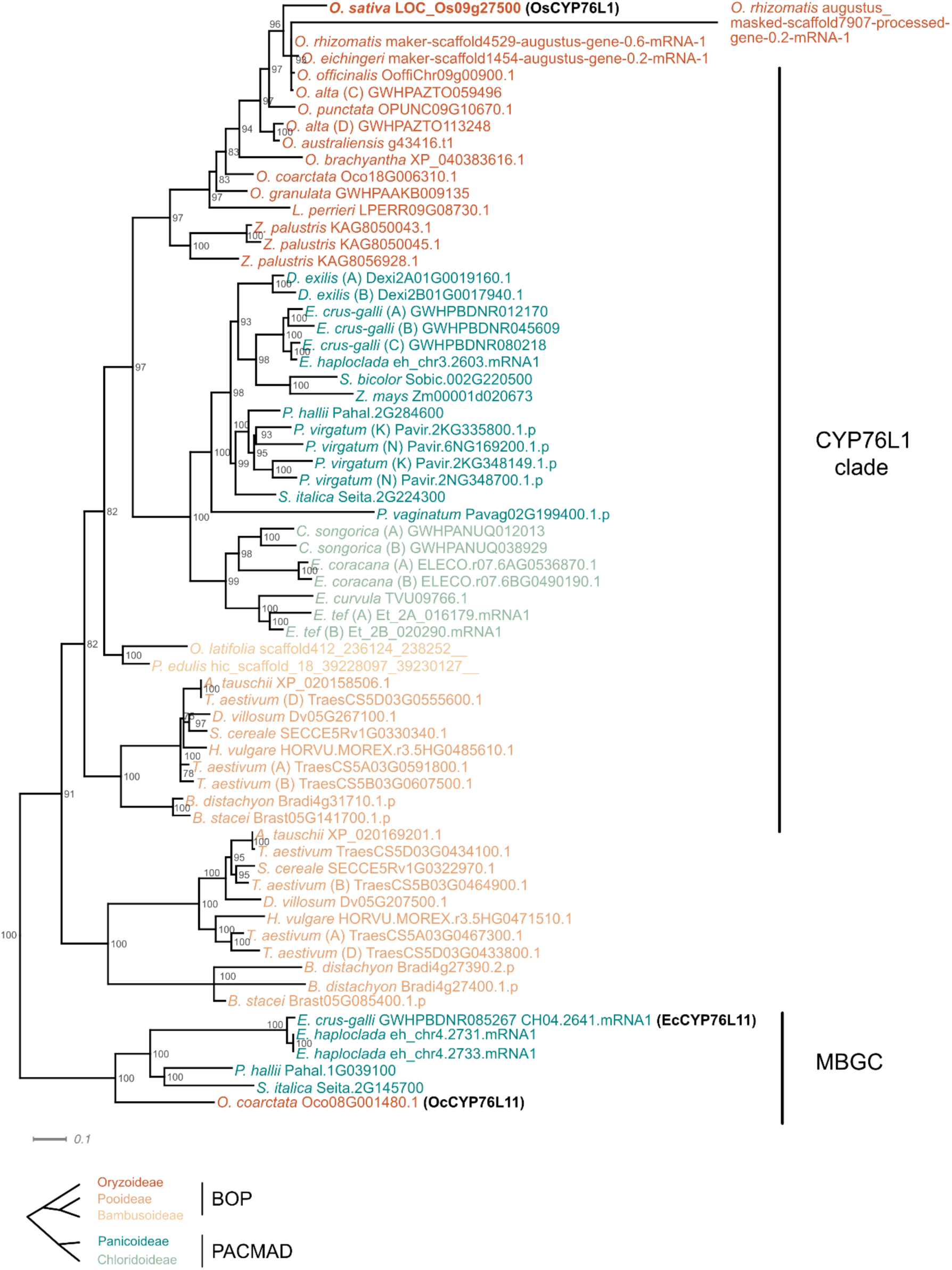
Phylogenetic analysis of a newly identified CYP ortholog in the MBGC. Subtree extracted from the maximum likelihood tree with the best-fit model selected by ModelFinder representing the relationship between the orthogroup containing *O. coarctata* CYP76L11 (see also Supplementary Figure 8). The phylogenetic tree is based on amino acid sequence alignments of the OsCYP76L1 clade, EcCYP76L11 and OcCYP76L11 (both present in the *E. crus-galli* and *O. coarctata* MBGC, respectively), and *Poaceae* species belonging to the subfamilies Oryzoideae, Pooideae, Bambusoideae, Panicoideae and Chloridoideae. The phylogenetic relation between these subfamilies is represented in the bottom graph.

### Momilactone production in different Oryza lineages

Among the investigated *Oryza* genome types (CC, CCDD, EE, KKLL and GG), only species and sub-genomes belonging to the lineages CC and LL include an orthologous MBGC. Notably, *O. coarctata* (KKLL), the earliest-divergent *Oryza* species with a MBGC, showed a distinct cluster architecture. To assess the functionality of the predicted clusters in the CC and LL lineages, we proceeded to evaluate the momilactone-producing potential of their respective representative species *O. officinalis* (CC) and *O. coarctata* (KKLL).

We conducted liquid chromatography-tandem mass spectrometry (LC-MS/MS) or ultra-high performance liquid chromatography (UHPLC)-MS/MS to detect momilactones A and B in leaf blades. Momilactone basal levels can be close to detection limits, which is why we induced momilactone production by treating the plants with CuCl_2_ 3 days before extraction, a treatment previously shown to increase momilactone levels in different *Oryza* species (Miyamoto et al., 2016). Consistent with the presence of the MBGC and the similar expression pattern of the corresponding genes upon induction (Supplementary Figure 9), we detected both momilactone A and B in *O. officinalis* (Figure 3a), but only momilactone A in *O. coarctata* (Figure 3b-e). Even after induction by CuCl_2_, *O. coarctata* accumulated only low levels of momilactone A (Figure 3b-e). Taken together, these results provide evidence for the functionality of the identified MBGC in the CC and KKLL species *O. officinalis* and *O. coarctata*, respectively.

**Figure 3.**
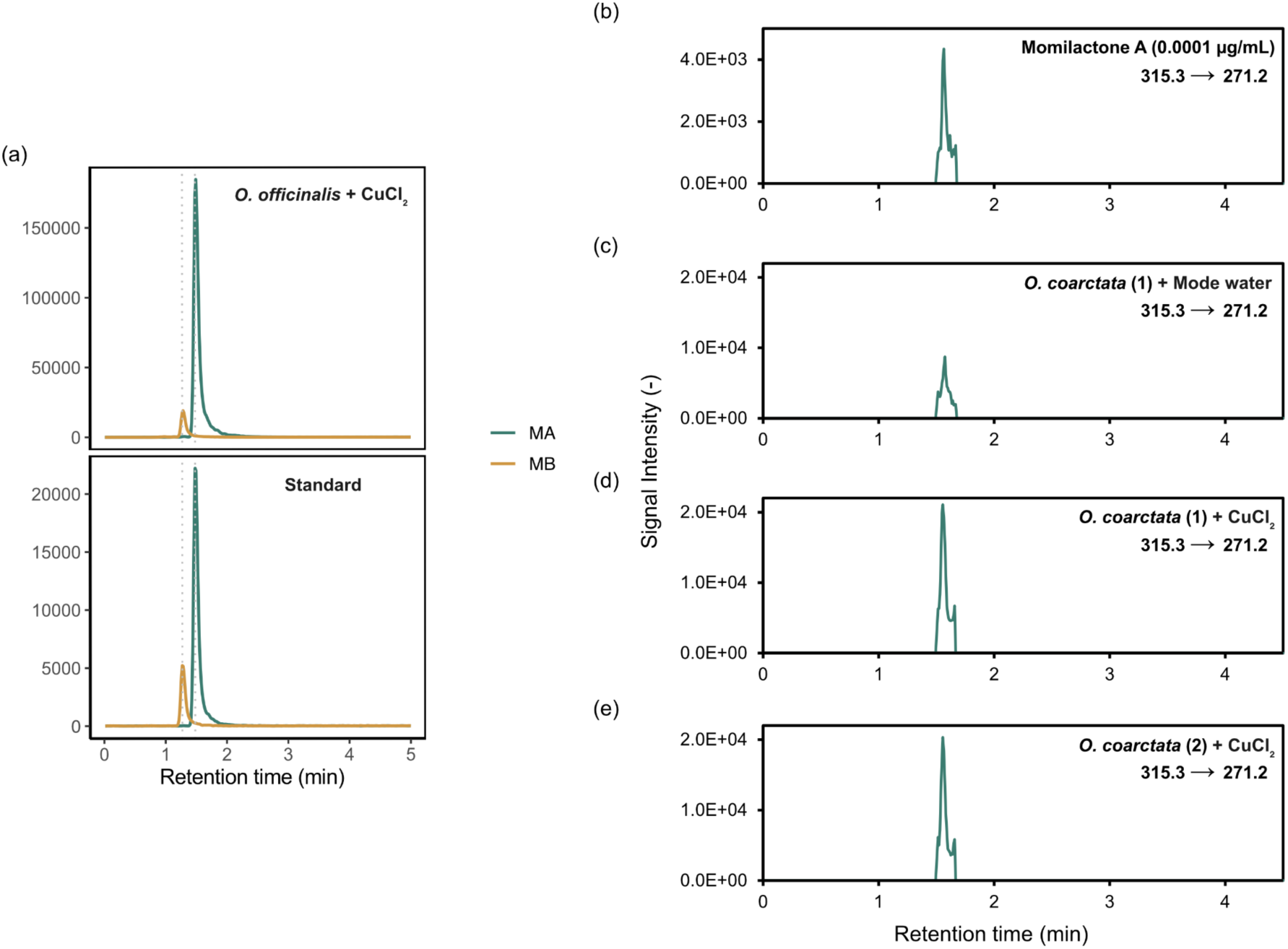
Analysis of momilactones in representative species of the *Oryza* genus. **(a)** Extracts from *O. officinalis* CuCl2-treated leaf blades were analysed by liquid chromatography-tandem mass spectrometry (LC-MS/MS). Five µl of the extract were subjected to LC-MS/MS under the conditions described in the Materials and Methods section. Momilactones were detected with combinations of m/z 315/271 for momilactone A and m/z 331/269 for momilactone B in the multiple reaction monitoring mode. MA, momilactone A; MB, momilactone **(b-e)**. Ultra-high performance liquid chromatography (UHPLC)-MS/MS (ESI^+^) chromatograms in the scheduled MRM mode of momilactone A in reference solution (0.0001µg/mL) **(b)**, mode water *O. coarctata* **(c)**, CuCl2-treated leaf blades of *O. coarctata* replicate 1 **(d)** and CuCl2-treated leaf blades of *O. coarctata* replicate 2 **(e)** showing the specific mass transition m/z 315.3 / 271.2 used for quantitation.

### Functional analysis of KSL4 in O. officinalis and O. coarctata

After establishing the existence of orthologous MBGCs in wild relatives of cultivated rice from the *Oryza* genus and confirming the presence of momilactone A and B in *O. officinalis* and momilactone A in *O. coarctata*, we next wanted to confirm the enzymatic functionality of the orthologs from those clusters. We focused on KSL4, which is known to catalyse the first dedicated step towards momilactone biosynthesis (*i.e.*, the conversion of *syn*-CDP to 9βH-pimara-7,15-diene). We tested the enzymatic activity of the KSL4 orthologs in representative species of the late-diverged CC and early-diverged KKLL lineages: OoKSL4-1 and OoKSL4-2 from *O. officinalis*, and OcKSL4 from *O. coarctata*.

The available annotation of the genes in the MBGC in *O. officinalis* showed inconsistencies, such as an extra sequence of amino acids in the C-terminal part of OoKSL4-1 and a missing exon in both *OoKSL4-1* and *OoKSL4-2*, compared to the ortholog in *O. sativa* (*OsKSL4*) (Supplementary Figure 10). These discrepancies may be attributed to limitations in the genome annotation process, which relied on RNA-seq data from tissues in which the constitutive expression of momilactone biosynthetic genes is generally low or absent (Supplementary Figure 9) (Shenton et al., 2020).

Exposure to UV light is known to induce the accumulation of momilactones in rice leaves (Kodama et al., 1988). To accurately investigate the function of KSL4 in *O. officinalis* and *O. coarctata*, we used previously published mRNA-Seq data from UV-irradiated leaf sheaths of both species to assemble a *de novo* transcriptome (Itoh et al., 2021). Subsequently, we conducted a homology search for OsKSL4 homologs in *O. officinalis* and *O. coarctata*. Using the assembled contig sequences as a reference, we designed specific primers and successfully cloned the cDNAs of these three *KSL4* homologs. We noticed that all cDNA clones of *OoKSL4-2*, obtained from independent plants of the same accession as the reference genome (W0002), exhibited a single nucleotide polymorphism (SNP) at position 7,882,960 that resulted in a G-to-A substitution, introducing a premature stop codon at amino acid position 445 (Supplementary Figure 10 and 11). We confirmed that all plants in our study were derived from one specific individual plant from the stock centre, and that this clone carried a *de novo* mutation that had been clonally propagated during plant material distribution; further investigation revealed that this SNP was absent in 14 other *O. officinalis* accessions, including the reference genome sequence of accession W0002, and hence is not representative of the consensus sequence at the population level (Supplementary Figure 11). We therefore reverted the G-to-A polymorphism in our cDNA clone to the consensus reference sequence of *OoKSL4-2*. We proceeded to confirm the enzymatic activity of OoKSL4-1, OoKSL4-2 and OcKSL4 by performing a *syn*-CDP conversion assay in *E. coli* and analysing the products using gas chromatography coupled to mass spectrometry (GC-MS). The retention time of the product generated by OoKSL4-2 and OcKSL4 was identical to that of 9βH-pimara-7,15-diene, indicating that both enzymes have the same enzymatic activity as OsKSL4 (Figure 4a). In contrast, we did not observe any production of 9βH-pimara-7,15-diene by OoKSL4-1, but production of minute amounts of an unidentified product (labelled as “b” in Figure 4a). We were able to corroborate the production of 9βH-pimara-7,15-diene by OoKSL4-2 by transiently expressing OoKSL4-1 and OoKSL4-2 in *Agrobacterium tumefaciens*-infiltrated *Nicotiana benthamiana* leaves (Figure 4b). We therefore conclude that OoKSL4-2 is the enzyme responsible for catalysing the conversion of *syn*-CDP to *syn*-pimaradiene in *O. officinalis*. Taken together, these results further support that the orthologous MBGC the we identified in *O. officinalis* and *O. coarctata* are functional and responsible for the production of momilactones.

**Figure 4.**
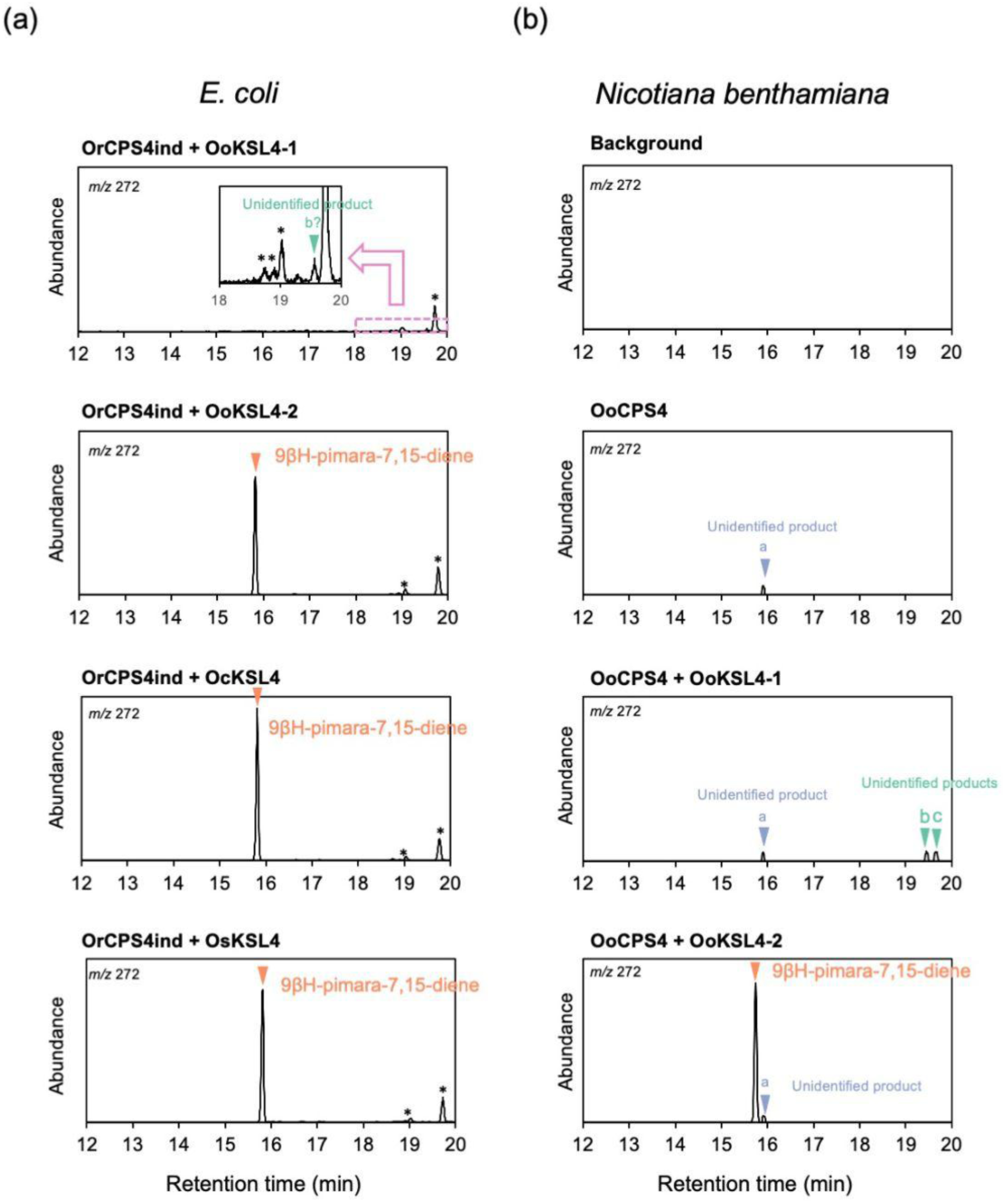
Functional characterization of KSL4 orthologs. **a)** GC-MS chromatograms of the products obtained from recombinant OoKSL4-1, OoKSL4-2 (with the reverted Trp at position 445), OcKSL4, and OsKSL4 from *E. coli* in a metabolic engineering system. **b)** GC-MS chromatograms of the products obtained in the *Nicotiana benthamiana* transient expression system after expressing *OoCPS4*, *OoCPS4* + *OoKSL4-1* and *OoCPS4* + *OoKSL4-2*. Orange triangles indicate production of 9βH-pimara-7,15-diene. Non-specific peaks from *E. coli* cells are indicated with asterisks (*). Mass spectra are included in Supplementary Figure 12.

## DISCUSSION

### The momilactone BGC shows diversified architecture within the Oryza genus

BGCs arise from the duplication of enzymes involved in primary metabolism, followed by neofunctionalization and/or relocation processes that are influenced by positive and negative selection pressures (Polturak et al., 2022b; Smit and Lichman, 2022). Ecological factors shape the direction and strength of the selection process, determining the evolution of the metabolism and ultimately affecting the architecture of BGCs. Here, we studied the composition and evolution of the momilactone biosynthetic gene cluster in previously unexplored lineages in the genus *Oryza*. We provide evidence that the MBGC is prevalent in the CC and LL lineages, while it is absent in the intermediate EE or the basal GG lineages (Figure 1b). Our data show that the evolution of the MBGC is marked by lineage-specific rearrangements, such as gene copy number variation or the increase of the size of the cluster, on-going pseudogenization in certain paralogs, and occasional cluster loss. Moreover, we found a distinct MBGC architecture in *O. coarctata*, with the cluster in that species harbouring a close ortholog of *EcCYP76L11*. This finding represents the first instance of an alternative architecture of the MBGC in *Oryza* and strengthens the idea of a common origin of the MBGC in *Echinochloa* and *Oryza*. Overall, our research sheds light on the evolutionary dynamics and distribution of the MBGC in the genus *Oryza*.

In our study, we found the MBGC at a basal position of the *Oryza* phylogeny in the species *O. coarctata* (sub-genome LL). *O. coarctata* is a polyploid species resulting from the hybridization of the KK and LL diploid genomes; the respective ancestral species have presumably gone extinct or remain to be discovered (Ge et al., 1999; Lu et al., 2009). The KK sub-genome occupies a basal position relative to the EE and DD lineages, while the *O. coarctata* LL sub-genome (initially designated as HH) occupies a more basal position compared to KK and it is closer to the diploid FF genome species *O. brachyantha*. Since we found the cluster in *O. coarctata*, a species that branched off before the divergence of the EE lineage, we hypothesize that *O. australiensis* (EE) likely lost the MBGC. Our study thereby is one of the very few to document a case of inter-specific loss of biosynthetic gene clusters (Smit and Lichman, 2022). A lineage-specific loss of a BGC was previously described in *Oryza* for the phytocassane biosynthetic gene cluster: *Oryza* AA species and the outgroup species *L. perrieri* and *Zizania palustris* possess the phytocassane cluster, whereas it was partially lost in *O. punctata* (BB) and *O. brachyantha* (FF) (Miyamoto et al., 2016; Yan et al., 2022). It is worth noting that the MBGC is also found in the *Panicoideae* and *Chloridoideae* subfamilies of the *Poaceae* and most likely evolved from a common ancestor (Wu et al., 2022a). However, the distribution of the cluster in the PACMAD clade is restricted to certain tribes, while it is either fragmented or completely lost in others (Wu et al., 2022a). In line with these observations, our results highlight the dynamic nature of biosynthetic gene clusters and suggest that loss of BGCs might be prevalent within *Poaceae*.

The MBGC and MBGC-like clusters are widespread in the *Poaceae* family (Wu et al., 2022a). Species belonging to the *Triticeae* tribe (Pooideae), phylogenetically more closely related to *Oryza* than to *Echinochloa*, possess a functionally distinct diterpene cluster that shares similarities with the MBGC (Liu et al., 2021; Polturak et al., 2022a; Wu et al., 2022a). This MBGC-like is located on the homeologous chromosomes 2A and 2D in wheat (*Triticum aestivum*), homologs of rice chromosome 4, and exhibits synteny with the region containing the MBGC in rice (Polturak et al., 2022a). However, the cluster in *Triticeae* lacks certain important momilactone biosynthesis genes, e.g. a *MAS* and a *syn-CPS* to produce *syn* labdane-related diterpenoids. Instead, the *Triticeae* cluster produces normal labdane-related diterpenoids in wheat (*Triticum aestivum*) and barley (*Hordeum vulgare*) (Liu et al., 2021; Polturak et al., 2022a). Owing to the shared synteny between the MBGC in rice and the MBGC-like in wheat, Polturak et al. suggested that the *Triticeae* diterpene cluster and the MBGC share a common evolutionary origin, and hypothesized that both clusters could have originated from a common ancestral BGC before the divergence of the PACMAD and BOP clades (Polturak et al., 2022a). Wu and colleagues, who performed a more extensive survey of the MBGC in different Poaceae genomes, alternatively attributed the evolution of the cluster in Poaceae to multiple instances of lateral gene transfer (LGT) and proposed that the MBGCs found in the PACMAD clade (Panicoideae and Chloridoideae subfamilies) originated from the MBGC-like cluster in *Triticeae* (Wu et al., 2022a). They moreover speculated that *EcCYP76L11* in *Echinochloa* was also laterally transferred from the *Triticeae* clade along with the MBGC-like. After the PACMAD species acquired the MBGC-like, it would then have recruited *MAS*. Subsequently, another LGT event would have had to occur, resulting in the acquisition of the cluster by *Oryzoideae*, followed by the loss of the *CYP76L11* gene (Wu et al., 2022a). Wu and colleagues based their conclusions on the phylogenetic incongruences among the MBGC orthologous genes, topology tests on the constrained trees for each MBGC gene, and a general lack of synteny between the genomic region containing the MBGC in the different tribes in the *Poaceae*. It is worth noting that the MBGC in *Panicoideae* and *Chloridoideae* is also not syntenic, following the same reasoning, this would imply that these gene clusters were acquired through two separate and independent lateral gene transfer (LGT) events rather than evolving from a common ancestor, as suggested by the same authors. There are several phylogenomic studies that suggest that LGT is prevalent among grasses; LGTs in *Poaceae* usually occur among closely related species, while the insertion occurs in random and non-syntenic genomic regions; however, the underlying mechanisms by which natural plant-plant LGT would occur are still largely unknown (Dunning et al., 2019; Hibdige et al., 2021). Our findings provide evidence that the *CYP76L11* gene has not been completely lost in the genus *Oryza*. The discovery of *CYP76L11* in *O. coarctata* implies either a common origin by vertical transmission of the MBGC between *Echinochloa* and *Oryza*, or a lateral gene transfer (LGT), as proposed by Wu et al. (2022). However, the findings from both Polturak et al. (2022a) and our study (Supplementary Figure 13) indicate high synteny between the flanking regions containing the MBGC-like cluster in *Triticeae* and the MBGC in *Oryza* (including *O. coarctata*) and thus suggest a common evolutionary origin of this genomic region, including the MBGC. This suggests that there may have been a common ancestor of the MBGC that existed before *Oryza* and *Triticeae* diverged. However, it is difficult to reconcile the synteny results shown by us and Polturak *et al*. (2022a) with the phylogenomic signatures of LGT shown by Wu et al., especially considering that the MBGC in *Echinochloa* and *Oryza* produces the same type of compounds, unlike in *Triticeae*. To shed light onto this question, it is important to consider the impact and prevalence of gene and MBGC losses in *Poaceae*. The presence or absence of MBGCs in different species can have significant implications for phylogenomic signals and evolutionary dynamics. The loss of MBGCs in certain lineages could potentially shape the phylogenomic patterns observed in Poaceae. Further investigations into the frequency and significance of MBGC losses across Poaceae will contribute to a better understanding of the evolutionary history of this cluster.

### Allotetraploid rice species carry only one MBGC copy in their genome

In our study, we aimed to investigate whether allotetraploid species retain two copies of the MBGC. Previous research has shown that whole genome duplication (WGD) can lead to the silencing and progressive loss of one copy of a biosynthetic gene cluster, likely to balance the effects of increased gene dosage (Polturak et al., 2022a; Yang et al., 2021). We found that each of the two allotetraploid species we studied had retained one copy of the MBGC. In *Oryza alta* (CCDD), the cluster was present in the CC but not the DD sub-genome, which means that the MBGC was either lost in the DD sub-genome after hybridization or that the DD ancestor had already lost it prior to the hybridisation. Given that *O. australiensis* (EE), the closest relative of the *O. alta* D-type sub-genome (the ancestral diploid D-type donor is presumably extinct), lacks a MBGC, it is possible that the cluster had already been lost in the DD donor.

Similarly, we identified only one copy of the MBGC in the *O. coarctata* LL but not the KK sub-genome. The LL sub-genome derives from a donor species that diverged earlier than the KK sub-genome and is more closely related to *O. brachyantha* (FF), a species that harbours an incomplete MBGC (Guo and Ge, 2005; Miyamoto et al., 2016). However, we cannot say with certainty whether the cluster was lost in the KK sub-genome after hybridization or was already lost prior to the hybridisation in the ancestor. Future studies using yet-to-be generated haplotype-resolved assemblies of BBCC hybrid species could contribute to a better understanding of cluster evolution in allotetraploid rice, as both ancestors of these hybrids harbour the MBGC.

### Momilactone B formation may be a recent innovation in the Oryza genus

At the interspecies level, the existence of conserved BGCs does not always align consistently with the production of identical metabolites, while the metabolites derived from orthologous BGCs can also be species-specific. The resulting divergence in chemotypes between species is associated with variations in biosynthetic genes external to the BGC, genes within the BGC, or transcription factors controlling the biosynthesis of these compounds (Liu et al., 2020; Zhou et al., 2016). In some cases, the function of the BGC is retained, and orthologous BGCs produce the same molecule. This is exemplified by *Oryza* species from the AA and BB lineages, which produce momilactones A and B (Miyamoto et al., 2016).

In our study, we show that the function of the MBGC is not only retained in the AA and BB but also the CC and LL lineages. Specifically, we found that the CC lineage, represented by *O. officinalis*, possesses an MBGC with slight differences in gene copy number, and that both momilactone A and B can be detected. The KKLL species *O. coarctata* contains an orthologous and syntenic MBGC with a distinct architecture compared to *O. sativa*. In line with our phylogenomic analysis suggesting the potential for momilactone production in *O. coarctata* and the co-expression of its MBGC genes, we were able to detect momilactone A in leaf blades induced with CuCl_2_ (Miyamoto et al., 2016). However, we did not detect momilactone B in *O. coarctata*. Among the auxiliary genes outside the MBGC, *CYP701A8* and *CYP76M8* are necessary for the production of momilactone A, while CYP76M14 catalyses the conversion of momilactone A to momilactone B (Figure 1a) (De La Peña and Sattely, 2021; Kitaoka et al., 2021). The lack of momilactone B in *O. coarctata* could be attributed to the suppressed expression of the *CYP76M14* ortholog (*Oc01G012750*), which was only weakly expressed (<3 TPM) upon UV treatment and barely detectable under control conditions in different tissues (Supplementary Figure 9). Intriguingly, unlike the MBGC genes, which are located on the LL sub-genome in *O. coarctata*, the only copy of *OcCYP76M14* is located on the KK sub-genome. To date, among the Poaceae, momilactone B has been detected only in AA, BB (Miyamoto et al. 2016) and CC *Oryza* species (this study). At a larger phylogenetic scale, *CYP76M14* is exclusive to *Oryza*, specifically to AA, BB, CC and KK lineages, and appears to have arisen from a duplication within the CYP76M subfamily (Supplementary Figure 8). It is imaginable that the production of momilactone B might have evolved after the divergence of the KK lineage by recruiting *CYP76M14* into the momilactone regulatory network.

## MATERIALS AND METHODS

### Oryza and Poaceae datasets

*O. sativa* (v7.0) (Ouyang et al., 2007) protein sequences and gene annotations were downloaded from Phytozome (https://phytozome-next.jgi.doe.gov/); *O. punctata* (Oryza_punctata_v1.2) (Stein et al., 2018) protein sequences and gene annotations from EnsemblPlants; the *O. officinalis* (Shenton et al., 2020) genome sequence from NCBI (assembly GCA_008326285.1) and annotations from the original article repository; *O. eichingeri* and *O. rhizomatis* (Shenton et al., 2020) protein sequences and gene annotations from Cyverse Data Commons (https://doi.org/10.25739/awh3-dm39); *O. alta* (Yu et al., 2021) protein sequences and gene annotations from the from the Genome Warehouse (GWH) accession number GWHAZTO00000000; the *O. australiensis* (Phillips et al., 2022) genome sequence from the National Center for Biotechnology Information (NCBI) (GCA_019925245.2) and annotations from figShare (https://figshare.com/collections/Oryza_australiensis_Keep_River_Genome_Assembly_V2/5 875592/2); the *O. coarctata* (Zhao et al., 2023) protein sequences and gene annotation from figShare (https://figshare.com/articles/dataset/A_high-quality_chromosome-level_wild_rice_genome_of_i_Oryza_coarctata_i_/23938590/1); the *O. brachyantha* (Chen et al., 2013) (ObraRS2) genome assembly and gene annotations from NCBI (GCF_000231095.2); *O. granulata* (Shi et al., 2020) protein sequences and gene annotations from GWH (GWHAAKB00000000); *L. perrieri* (Stein et al., 2018) protein sequences and gene annotations (V1.4) from EnsemblPlants*. O. officinalis*, *O. australiensis* and *O. brachyantha* protein sequences were extracted from the gene annotation using gffread (v0.12.7) (Pertea and Pertea, 2020).

*Aegilops tauschii subsp. strangulata* (V5.0; GCF_002575655.2) (Wang et al., 2021) protein sequences were retrieved from NCBI, *Brachypodium stacei* (v1.1) (Gordon et al., 2020) from Phytozome, *Brachypodium distachyon* (V3.1) (Sreedasyam et al., 2023; Vogel et al., 2010) from Phytozome; *Cleistogenes songorica* (Zhang et al., 2021) from GWH accession PRJCA002752; *Digitaria exilis* (CM05836) (Abrouk et al., 2020) from article repository in Dryad; *Eleusine coracana* (v1.1) (Devos et al., 2023) from Phytozome; *Echinochloa crus-galli* (V3.0) (Wu et al., 2022b) sequences from GWH, accession number GWHBDNR00000000; Echinochloa haploclada from (Ye et al., 2020) article repository; *Eragrostis curvula* (CERZOS_EC1.0; GCA_007726485.1) (Carballo et al., 2019) from NCBI; *Eragrostis tef* (ASM97063v1) (Cannarozzi et al., 2014) from Ensembl Plants; *Hordeum vulgare* cv Morex (V3) (Mascher et al., 2021) sequences from e!DAL (DOI: 10.5447/ipk/2021/3), we specifically used the file Hv_Morex.pgsb.Jul2020.aa.fa that includes low confidence proteins. Unlike in databases such as Phytozome and EnsemblPlants, this annotation includes most of the gene in the diterpene cluster reported by (Liu et al., 2021); *Olyra latifolia* (Guo et al., 2019) sequences from http://www.genobank.org/bamboo; *Oropetium thomaeum* (v1.0) (VanBuren et al., 2015) from Phytozome; *Phyllostachys edulis* (Zhao et al., 2018) from http://gigadb.org/dataset/100498; *Panicum hallii* var. *filipes* (V3.2) (Lovell et al., 2018) sequences from Phytozome; *Panicum virgatum* (v5.1) (Lovell et al., 2021; Sreedasyam et al., 2023) from Phytozome; *Paspalum vaginatum* (v3.1) (Sun et al., 2022) from Phytozome; *Pharus latifolius* (Ma et al., 2021) sequences from http://www.genobank.org/grass; *Sorghum bicolor* (V3.1.1) (McCormick et al., 2018) from Phytozome, *Secale cereale* (Lo7_2018v1p1p1.pgsb.Feb2019.HC) (Rabanus-Wallace et al., 2021) from article repository; *Setaria italica* (V2.2) (Bennetzen et al., 2012) sequences from Phytozome; *Triticum aestivum* cv Chinese Spring (IWGSC V2.1) (Zhu et al., 2021) from https://urgi.versailles.inrae.fr/download/iwgsc/IWGSC_RefSeq_Annotations/v2.1/ (file iwgsc_refseqv2.1_annotation_200916_HC_pep.valid), *Zea mays* B73 (Schnable et al., 2009) (V4) from Phytozome and *Zizania palustris* (Haas et al., 2021) from NCBI (accession JAAALK000000000) and *Ananas comosus* (V3) (Ming et al., 2015) proteome from Phytozome as outgroup.

### MBGC ortholog detection

OrthoFinder (v2.5.4) (Emms and Kelly, 2019, 2015) was used to infer orthogroups containing *O. sativa* momilactone biosynthesis genes (i.e., genes from multiple species descended from a single gene in the last common ancestor of these species). In the case of the *Oryza* dataset, the input for OrthoFinder included primary transcript proteomes of the following species: *O. sativa*, *O. punctata*, *O. officinalis*, *O. eichingeri*, *O. rhizomatis*, *O. alta*, *O. australiensis*, *O. coarctata*, *O. brachyantha*, *O. granulata*, and *L. perrieri* (included as the out-group). We used the ‘-M msa’ option to also infer the species tree (described in Emms and Kelly, 2018). Briefly, a maximum-likelihood species tree was inferred by using concatenated multiple sequence alignment of single-copy orthologs using 4069 orthogroups with a minimum of 100.0% of species having single-copy genes in any orthogroup.

Each orthogroup was aligned using MAFFT (v7.490) (Katoh et al., 2002; Katoh and Standley, 2013). IQTREE (v2.1.4-beta) (Minh et al., 2020) was used to infer the maximum likelihood (ML) tree with the best-fit model selected by ModelFinder (Kalyaanamoorthy et al., 2017) with 1000 replicates of ultrafast bootstrapping (Hoang et al., 2018). ML rooted trees of each orthogroup were used to identify orthologs of *O. sativa* momilactone biosynthetic genes in *Oryza*. Genes within the same clade as *O. sativa* momilactone biosynthesis genes were considered as potentially clustered homologs. The genomic location upstream and downstream of these homologs was further analyzed to confirm clustering. Trees were represented with Dendroscope (v3.8.2).

As a quality control measure to ensure no potential momilactone genes were missed, we performed nBLAST using the genomic locus of each momilactone gene from *O. sativa* against each of the *Oryza* species studied. Where necessary, we manually annotated the given locus (i. e. *OoCYP76M14*).

### Oryza coarctata CYP76L11 orthologs

To infer the relationship between *EcCYP76L11 and OcoCYP76L11*, we used OrthoFinder to infer the orthogroup containing OcoCYP76L11. The input for OrthoFinder included the primary transcript proteomes of the *Oryza* species described previously, together with *Z. palustris* (Oryzoideae), *T. aestivum*, *A. tauschii*, *S. cereale*, *D. villosum*, *H. vulgare*, *B. distachyon*, *B. stacei* (Pooideae); *P. edulis*, *O. latifolia* (Bambusoideae); *Z. mays*, *S. bicolor*, *P. vaginatum*, *E. crus-galli*, *E. haploclada*, *S. italica*, *P. hallii*, *P. virgatum*, *D. exilis* (Panicoideae); *E. coracana*, *C. songorica*, *O. thomaeum*, *E. curvula*, *E. tef* (Chloridoideae); *P. latifolius*, as outgroup to the core Poaceae; and *A. comosus* as outgroup of Poaceae.

The orthogroup containing OcoCYP76L11 was aligned using MAFFT (v7.490) followed by the inference of the maximum likelihood (ML) tree with IQTREE (v2.1.4-beta) and selecting the best-fit model by ModelFinder with 1000 replicates of ultrafast bootstrapping.

### Microsynteny Analysis

Microsynteny analysis and figures were done with MCScan implemented in Python (v3.9) with JCVI utility libraries (v1.1.11) following the package workflow: https://github.com/tanghaibao/jcvi/wiki/MCscan-(Python-version). Scripts are detailed in https://github.com/spriego/Priego-Cubero-et-al.-Oryza. Briefly, protein FASTA files and GFF3 files of each species were used as input. When needed, the FASTA files were processed using Biopython’s SeqIO package (Cock et al., 2009). Pairwise ortholog and synteny blocks search was performed with default settings using a lift-over distance of 5, a c-score = 0.99 and a maximum of 1 iterations. Multi-synteny blocks were constructed combining the syntenic blocks detected on every species with respect to *Oryza sativa*.

### *RNA-seq* mapping and quantification

Publicly available RNA-seq datasets from *O. officinalis* and *O. coarctata* were used (see Supplementary Figure 9). RNA-seq mapping and quantification was done using the nf-core pipeline rnaseq v3.14 (https://nf-co.re/rnaseq/3.14.0) in Nextflow 22.10.4 (Patel et al., 2024). Briefly, adapter and quality trimming was performed using TrimGalore v0.6.7 (Krueger et al., 2021). Then, reads were mapped to reference genomes using STAR v2.7.9a (Dobin et al., 2013) and quantified using salmon v1.10.1 (Patro et al., 2017). Finally, feature counts were normalised by gene length and sequencing depth using transcripts per million (TPM), that were used to inspect the expression of the momilactones biosynthetic genes.

### SNP calling

Short-read Illumina sequencing data for 15 *O. officinalis* accession were downloaded from OryzaGenome 2.1 (http://viewer.shigen.info/oryzagenome21detail/index.xhtml) (Kajiya-Kanegae et al., 2021). SNP calling was performed using the nf-core pipeline sarek v3.0.1 with default parameters in Nextflow 22.10.4 (Cannarozzi et al., 2014; Ewels et al., 2020; Garcia et al., 2020). Briefly, the fastq files underwent sequence quality assessment and pre-processing using fastp v0.23.2 (Chen et al., 2018). The reads were mapped to the *O. officinalis* reference genome using BWA mem v0.7.17-r1188 (Katoh and Standley, 2013). Duplicates were marked with GATK MarkDuplicates version 4.2.6.1 (McKenna et al., 2010). Variant calling was performed with HaplotypeCaller (Poplin et al., 2018). The BaseRecalibrator step was skipped as it requires a database of known polymorphic sites, which is unavailable for *O. officinalis.* The output vcf files were merged with bcftools v1.13 (Danecek et al., 2021) and SNPs effect were annotated with snpEff v5.0.1 (Cingolani et al., 2012).

### Data analysis

RStudio with R version 4.3.1 was used for generating Figures 1b, 3a, and Supplementary figures 9 and 11. Scripts used for data analysis are available at https://github.com/spriego/Priego-Cubero-et-al.-Oryza. To generate the figures, the following R packages were used: Tidyverse (Wickham et al., 2019), khroma (Frerebeau, 2023), ggrepel https://github.com/slowkow/ggrepel, readxl https://github.com/tidyverse/readxl, MetBrewer (https://github.com/BlakeRMills/MetBrewer), and MoMAColors (https://github.com/BlakeRMills/MoMAColors).

### Plant materials

Seeds of *O. officinalis* (accession W0002) and *O. coarctata* (accession W0551), and genome DNA solutions of several accession of *O. officinalis* (W0002, W0566, 0614, W1131, W1200 and W1252) were provided by the Resource Bank of the National Institute of Genetics, Japan (http://www.shigen.nig.ac.jp/rice/oryzabase/locale/change).

### Momilactone measurements

Each harvested leaf blade was treated with 0.5 mM CuCl_2_ for 72 h before the analysis. The treated sample (roughly 100 to 500 mg) was submerged in 4 ml of 70% methanol at 4°C for 24 h.

For *O. officinalis*, 2 uL of the extract was subjected to LC-MS/MS analysis (Sciex QTRAP3200) with the following selected reaction monitoring transitions used as previously described (Miyamoto et al., 2016): momilactone A, m/z 315/271; momilactone B, m/z 331/269.

For *O. coarctata*, plant material (100-150 mg) was weighed in a 2 mL extraction tube (CKMix, Bertin Technologies, Montigny le Bretonneux, France), and the tube filled up with methanol/water (70/30, v/v, 2 mL). After extractive grinding (3 × 30 s with 30 s breaks, 6000 rpm) using the bead beater (Precellys Homogenizer, Bertin Technologies, Montigny le Bretonneux, France), samples were incubated for 24 h at 4°C. After centrifugation, the supernatant was evaporated to dryness, resumed in acetonitrile (50 µL), and injected into the UHPLC−MS/MS-system (5 µL). A QTRAP 6500+ mass spectrometer (Sciex, Darmstadt, Germany) was used to acquire mass spectra and product ion spectra in positive electrospray ionization (ESI) mode. The MS/MS system was operated in the scheduled multiple reaction monitoring (MRM) mode at an ion spray voltage of 5500 V with the following ion source parameters: curtain gas (35 psi), temperature (550°C), gas 1 (55 psi), gas 2 (65 psi), and entrance potential (10 V). For analysis of momilactones A and B, the MS/MS parameters were tuned to achieve fragmentation of the [M+H]+ molecular ions into specific product ions: [M+H]+: momilactone A m/z 315.2/271.2 (quantifier) and m/z 315.2/269.2 (qualifier), momilactone B m/z 331.3/269.2 (quantifier) and m/z 331.2/128.1 (qualifier). For tuning, acetonitrile/water solutions of each analyte were introduced by means of flow injection using a syringe pump. The samples were separated by means of an ExionLC UHPLC (Shimadzu Europa GmbH, Duisburg, Germany) consisting of two LC pump systems ExionLC AD, an ExionLC degasser, an ExionLC AD autosampler, an ExionLC AC column oven and an ExionLC controller, and equipped with a BEH Premier C18 column (150 × 2.1 mm, 100 Å, 1.7 μm, Waters, Manchester, UK). Operated with a flow rate of 0.5 mL/min at a column oven temperature of 45°C using 0.1% formic acid in water (v/v) as solvent A and 0.1% formic acid in acetonitrile (v/v) as solvent B. Chromatography was performed with the following gradient: 70% B increased to 77.5% in 2.5 min, increased to 100% B in 0.5 min, held 1 min at 100% B, decreased to 70% B in 0.5 min, and held 0.5 min at 70% B. Data acquisition, and instrumental control was performed with Analyst 1.6.3 software (Sciex, Darmstadt, Germany).

### cDNA cloning of OoKSL4 and OcKSL4 and their functional analysis by metabolic engineering systems

The nucleotide sequences of full-length *OoKSL4-1*, *OoKSL4-2* and *OcKSL4* have been deposited in the GenBank database with accession numbers LC831403, LC831404 and LC831406, respectively.

For *OoCPS4*, full length of cDNA was amplified by RT-PCR using cDNA prepared from RNA sample of *O. officinalis* leaf 24-hour after the CuCl_2_ treatment with forward primer (OoCPS4F: 5’-TTCTGCCCAAATTCGATGCCGGCCTTTACTGCATC-3’), reverse primer (OoCPS4R: 5’-GTGATGGTGATGCCCAATCACATCTTGGAATATGA-3’). Nucleotide sequence of OoCPS4 has been deposited in the GenBank database with accession number LC831405. For *OoKSL4-1* full length cDNA were amplified by RT-PCR using forward primer (OoKSL4-1F: 5’-TTCTGCCCAAATTCGATGGTTATCACCCATATTTTGAG-3’), reverse primer (OoKSL4-1R: 5’-GTGATGGTGATGCCCAGATGAACTTGTAGCACCTAG-3’). To recover the nonsense G-to-A polymorphism in *OoKSL4-2*, a 2-step overlap PCR was performed. For the first amplicon, forward primer-1 (OoKSL4-2F1: 5’-TTCTGCCCAAATTCGATGGCGAATCCTATGG-3’) and reverse primer-1 (OoKSL4-2R1: 5’-GCTACCTGACCAACAACCCATTT-3’) were used. For the second amplicon, forward primer-2 (OoKSL4-2F2: 5’-AAATGGGTTGTTGGTCAGGTAGC-3’) and reverse primer-2 (OoKSL4-2R2: 5’-GTGATGGTGATGCCCGTAGCCTATAGTTTTCAG-3’) were used. The full length of the recovered *OoKSL4-2* was then amplified using forward primer-1 and reverse primer-2 as the primer set, with amplicons from part 1 and part 2 serving as templates. For *OsDXS3* full length cDNA were amplified by RT-PCR using forward primer (DXS3F: 5’-TTCTGCCCAAATTCGATGGCGCTCCAGGCATCG-3’), reverse primer (DXS3R: 5’-GATGCATACCGGTCGTCAGCTGAGCTGAAGTGC-3’). For *OsGGPPS* full length cDNA were amplified by RT-PCR using forward primer (GGPSF: 5’-TTCTGCCCAAATTCGATGGCTGCCTTCCCCCCG-3’), reverse primer (GGPSR: 5’-GATGCATACCGGTCGTCAGTTCTGCCGATAGGC-3’).

Full length cDNAs of *OoCPS4*, *OoKSL4-1* and *OoKSL4-2*, containing transit peptides, and two genes from *O. sativa* encoding OsDXS3 and OsGGPPS in the MEP pathway were respectively cloned into NruI-SmaI sites of pEAQ-HT vector by In-Fusion cloning system (Takarabio-Clontech). Constructed pEAQ-HT-*OoffKSL4-1*, pEAQ-HT-*OoffKSL4-2*, pEAQ-HT-*OsDXS3* and pEAQ-HT-*OsGGPPS*, all with 6×His tags at the C-terminal region, were then transferred into *Agrobacterium tumefaciens* for agroinfiltration.

In the *Nicotiana benthamiana* transient expression system, 5-week-old plants grown in a plant incubator (23°C, 16 h light/8 h dark photoperiod) were used. After agroinfiltration, the plants were further grown for five days under the same conditions. The leaves were then harvested, cut into 5 mm squares, immersed in *n*-hexane for a one-day extraction. Extraction and purification were done as described previously (Toyomasu et al., 2018).

The cDNA encoding partial OoKSL4-1 for functional analyses using *E. coli* was amplified by RT-PCR using primers 5’-<u>GGATCC</u>ATGGCGAATCCTGTGGAAG-3’ (forward; BamHI site is underlined) and 5’-<u>GTCGAC</u>TTAAGATGAACTTGTAGCACCTAG-3’ (reverse; SalI site is underlined), and subcloned into pGEX-4T-3 (GE Healthcare) using BamHI and SalI. The cDNA encoding pseudo-mature OoKSL4-2 was amplified by RT-PCR using forward primer 5’-<u>GGTTCCGCGTGGATCC</u>GAGGCTAGAATAAGGAGGC-3’ and reverse primer 5’-<u>GGGAATTCGGGGATCC</u>CTAGTAGCCTATAGTTTTC-3’ (terminal sequences of pGEX-4T-3 digested by BamHI are underlined), and inserted into BamHI-digested pGEX-4T-3 by using In-Fusion Smart Assembly Cloning Kit (Takarabio-Clontech). Full-length *OcKSL4* cDNA was amplified by RT-PCR using primers 5’-<u>GGATCC</u>ATGGCGAATTATCCCATGGAG-3’ (forward; BamHI site is underlined) and 5’-TTAAGATGGACTAGTAGCCTCTAGTTTAAGTGGC-3’ (reverse; NotI site is underlined), and subcloned into pGEM-T-Easy. *OcKSL4* cDNA was inserted into pGEX-4T-3 using BamHI and NotI. For RT-PCR, we used cDNA prepared from RNA from leaf blades of *O. officinalis* W0002 and *O. coarctata* W0551 24 h after UV treatment. OoKSL4-2_445W was generated by mutagenesis PCR using OoKSL4-2 in pGEX as template, with primers 5’-gTCAGGTAGCTTATTGAAAC-3’ (forward) and 5’-caACAACCCATTTTCTCTAGGATTG-3’ (reverse; small characters indicate W codon for mutagenesis).

The functional analyses of OoKSL4-1, OoKSL4-2 and OcKSL4 in a metabolic engineering system using *E. coli* cells were performed as described previously (Toyomasu et al., 2016). The *n*-hexane extract from culture medium containing recombinant *E. coli* BL21-DE3 harbouring pRSF-PaGGS:OrCPS4ind and a respective construct to express one of the above proteins was subjected to GC-MS analysis. PaGGS is a geranylgeranyl synthase domain of fusicoccadiene synthase in *Phomopsis amygdali* (PaFS) (Toyomasu et al., 2007) and OrCPS4ind is a *syn*-CDP synthase from *O. rufipogon* W0106 (Miyamoto et al., 2016), to supply GGDP and *syn*-CDP, respectively.

GC-MS was conducted using an Agilent 8860 GC-5977 MSD system, fitted with a fused silica, chemically bonded capillary column (HP-1MS; 0.25 mm in diameter, 60 m long, 0.25 μm film thickness). Ionization voltage was 70 eV. Each sample was injected onto the column at 250°C in the splitless mode. After a 2-min isothermal hold at 60℃, the column temperature was increased by 30℃/min to 150℃, 10℃/min to 180℃, 2℃/min to 210℃, and 30℃/min to 300℃. The flow rate of the helium carrier gas was 1 mL/min. We used 9βH-pimara-7,15-diene as a standard, which was synthesized as previously described (Ye et al., 2017).

## AUTHOR CONTRIBUTIONS

S.P.C. and C.B. conceived the study. S.P.C. carried out the genomic and phylogenetic analyses. T.T. and Y.H. carried out in vitro enzymatic assays. the functional analysis of KSL4. Y.L. and K.O. carried out momilactone measurements. H.N., K.O., and C.B. supervised the work. S.P.C. and C.B. wrote the manuscript with contributions from all authors.

## ACKNOWLEDGEMENTS

We are grateful to José Villaécija Aguilar and Niklas Schandry for critical reading of the manuscript and to Niklas Schandry for setting up and maintaining environments for computational analyses. We would like to thank Reshi Shanmuganathan for help with identifying orthologs. We acknowledge the Graduate School Life Science Munich (LSM) for their valuable support to S.P.C. We would also like to thank Prof. Yutaka Sato and Mr. Hiroyasu Furumi at the National Institute of Genetics, Japan, for providing wild rice material, and Chisato Goda (Yamagata University, Japan) for technical support.

## FUNDING

This work was supported by the Deutsche Forschungsgemeinschaft (DFG; grant no. 457739273 (C.B.) and by JSPS KAKENHI Grant Numbers 20H02922 (K.O.) and 23H02148 (T.T.). For computational analyses, we used the BioHPC-Genomics compute cluster at the Leibniz Rechenzentrum (DFG; grant no. 450674345).

## SUPPLEMENTARY MATERIAL

### SUPPLEMENTARY FIGURES

**Supplementary Figure 1.**
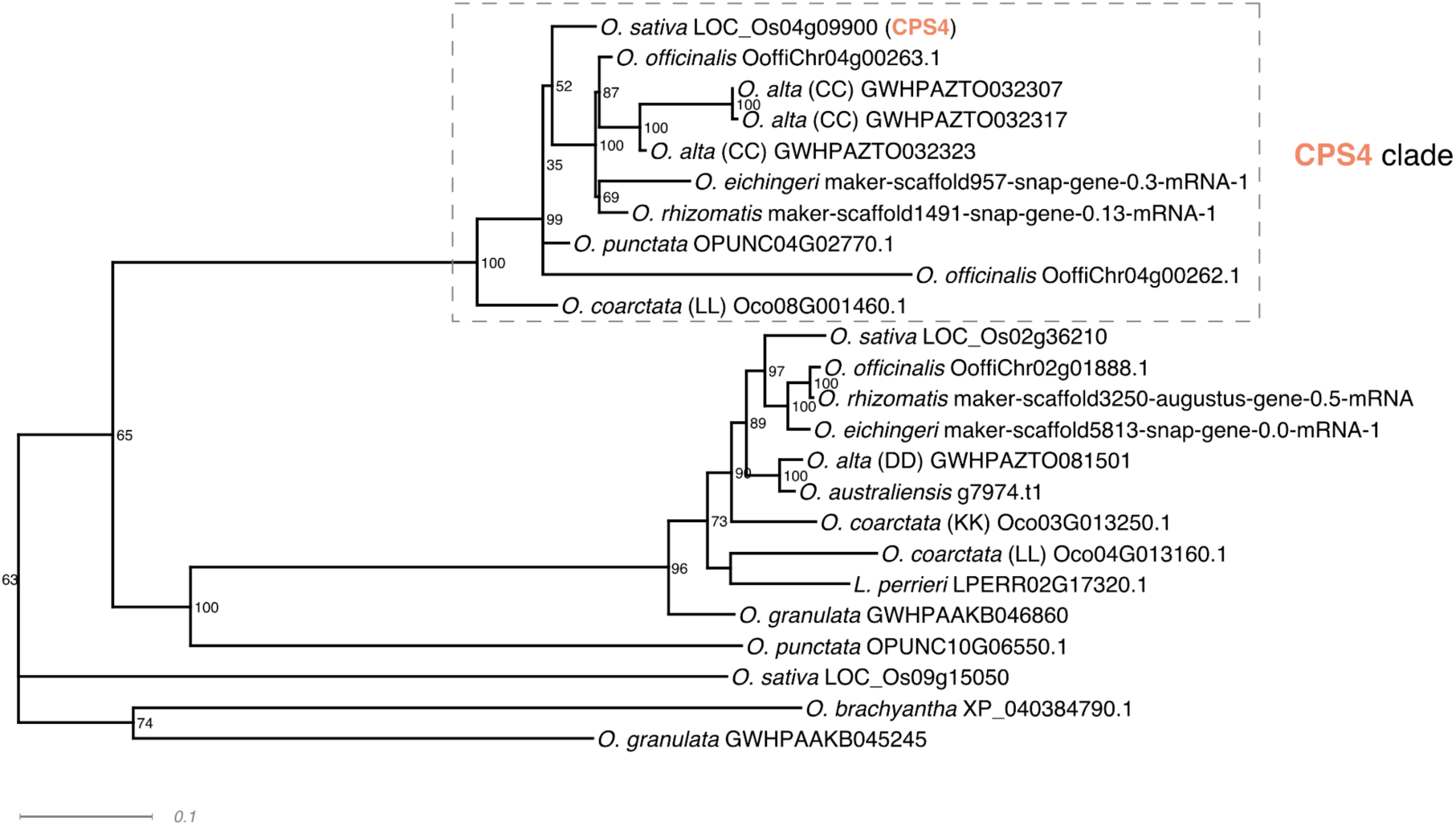
Phylogenetic analysis based on CPS4 amino acid sequence. Maximum likelihood tree with the best-fit model selected by ModelFinder for the orthogroup containing *O. sativa* CPS4 (LOC_Os04g09900) in *O. sativa*, *O. punctata*, *O. officinalis*, *O. eichingeri*, *O. rhizomatis*, *O. alta*, *O. australiensis*, *O. coarctata*, *O. brachyantha*, *O. granulata* and *L. perrieri*. Numbers represent 1000 replicates of ultrafast bootstrapping. Dashed rectangle highlights the CPS4 clade, which contains orthologs potentially involved in momilactones biosynthesis and clustered in *Oryza* species.

**Supplementary Figure 2.**
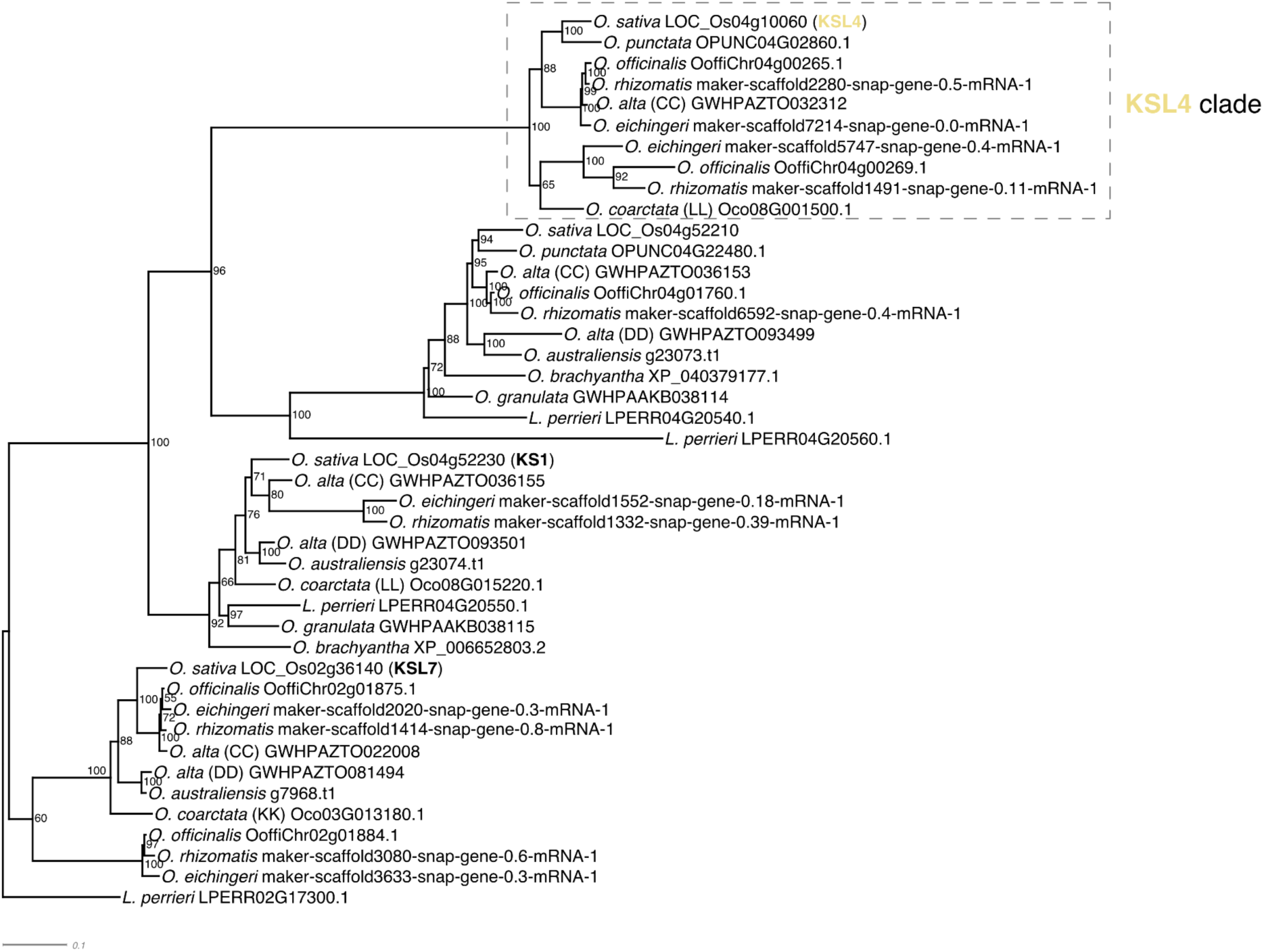
Phylogenetic analysis based on KSL4 amino acid sequence. Maximum likelihood tree with the best-fit model selected by ModelFinder for the orthogroup containing *O. sativa* KSL4 (LOC_Os04g10060) in *O. sativa*, *O. punctata*, *O. officinalis*, *O. eichingeri*, *O. rhizomatis*, *O. alta*, *O. australiensis*, *O. coarctata*, *O. brachyantha*, *O. granulata* and *L. perrieri*. Numbers represent 1000 replicates of ultrafast bootstrapping. Dashed rectangle highlights the KSL4 clade, which contains orthologs potentially involved in momilactones biosynthesis and clustered in *Oryza* species.

**Supplementary Figure 3.**
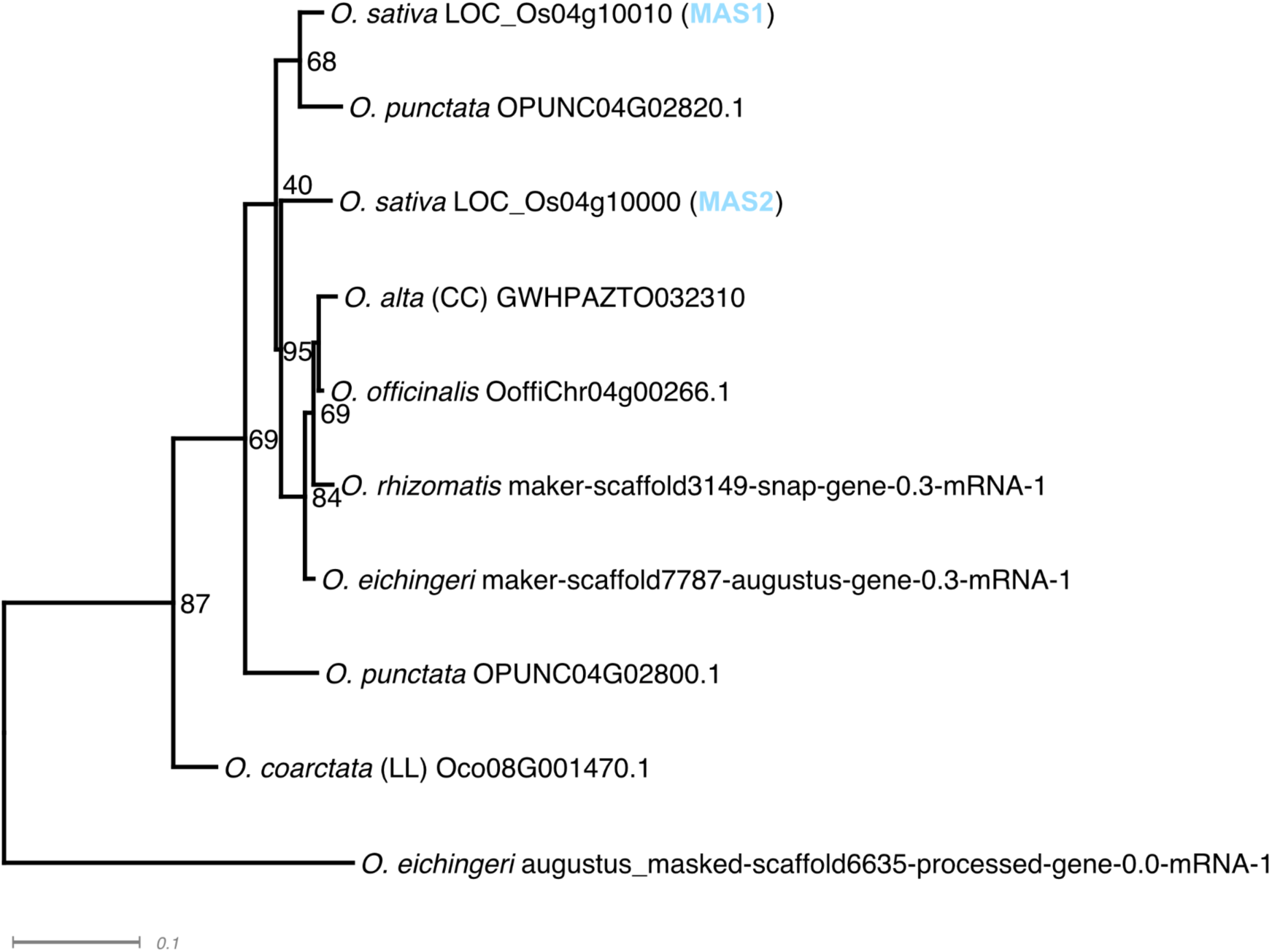
Phylogenetic analysis based on MAS amino acid sequences. Maximum likelihood tree with the best-fit model selected by ModelFinder for the orthogroup containing *O. sativa* MAS1 and MAS2 (LOC_Os04g10010, LOC_Os04g10000) in *O. sativa*, *O. punctata*, *O. officinalis*, *O. eichingeri*, *O. rhizomatis*, *O. alta*, *O. australiensis*, *O. coarctata*, *O. brachyantha*, *O. granulata* and *L. perrieri*. Numbers represent 1000 replicates of ultrafast bootstrapping. Since this orthogroup consisted of a small number of genes, we considered all of them as potentially involved in momilactone biosynthesis and clustered.

**Supplementary Figure 4.**
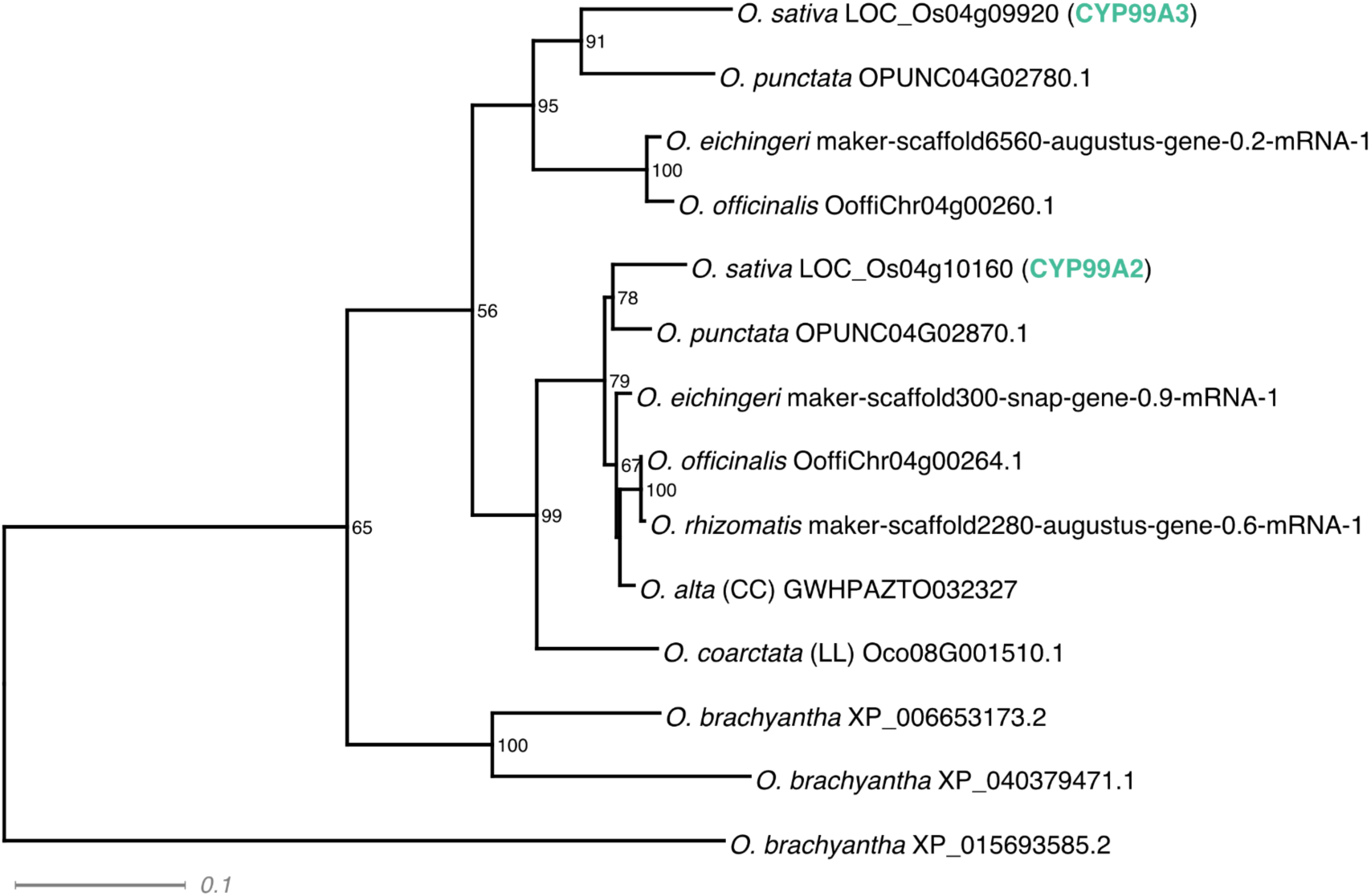
Phylogenetic analysis based on CYP99A amino acid sequences. Maximum likelihood tree with the best-fit model selected by ModelFinder for the orthogroup containing *O. sativa* CYP99A2 and CYP99A3 (LOC_Os04g10160, LOC_Os04g09920) in *O. sativa*, *O. punctata*, *O. officinalis*, *O. eichingeri*, *O. rhizomatis*, *O. alta*, *O. australiensis*, *O. coarctata*, *O. brachyantha*, *O. granulata* and *L. perrieri*. Numbers represent 1000 replicates of ultrafast bootstrapping. Since this orthogroup consisted of a small number of genes, we considered all of them as potentially involved in momilactone biosynthesis and clustered.

**Supplementary Figure 5.**
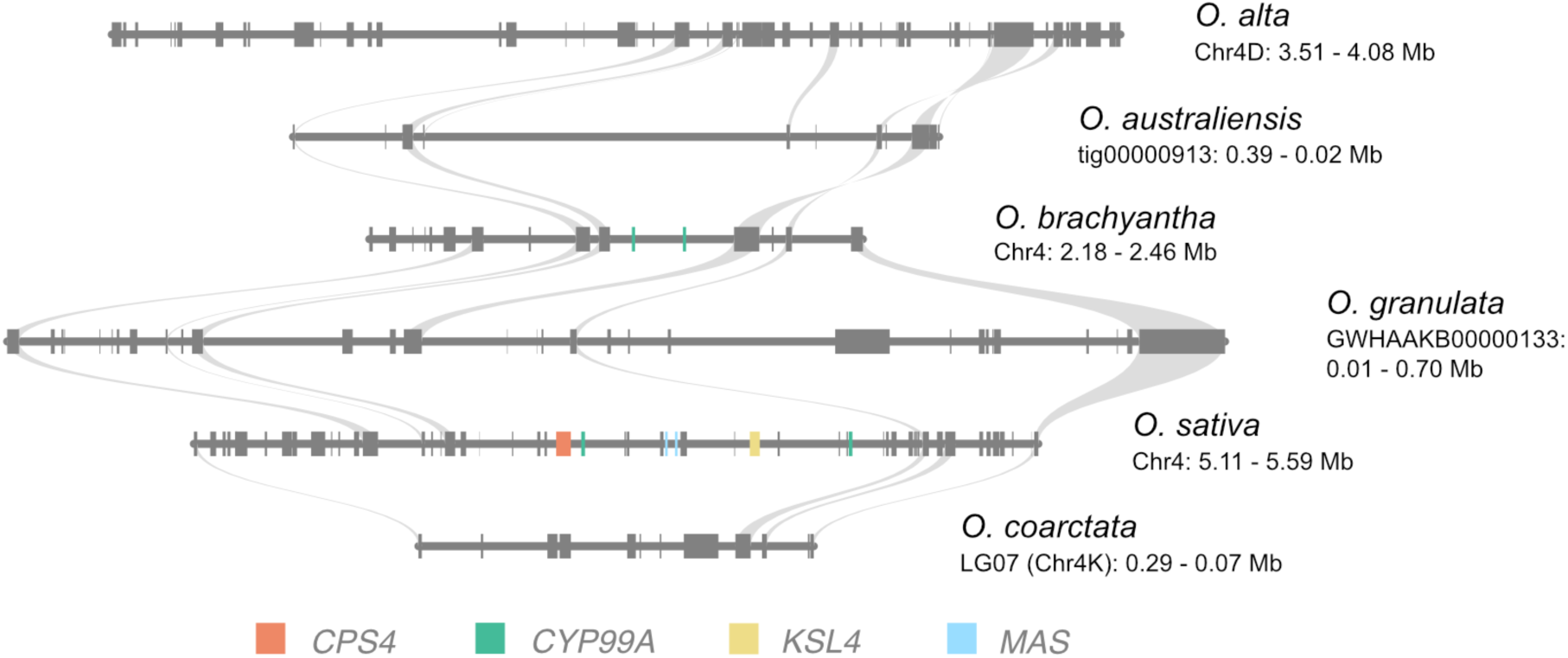
Microsynteny of the MBGC between *O. sativa* and species and sub-genomes lacking a MBGC. *O. alta* sub-genome DD was used as anchor species. The region was not further expanded given the fragmented nature of the assemblies for *O. australiensis* and *O. granulata*. Coloured rectangles represent the momilactone biosynthetic genes. Grey lines connect orthologs.

**Supplementary Figure 6.**
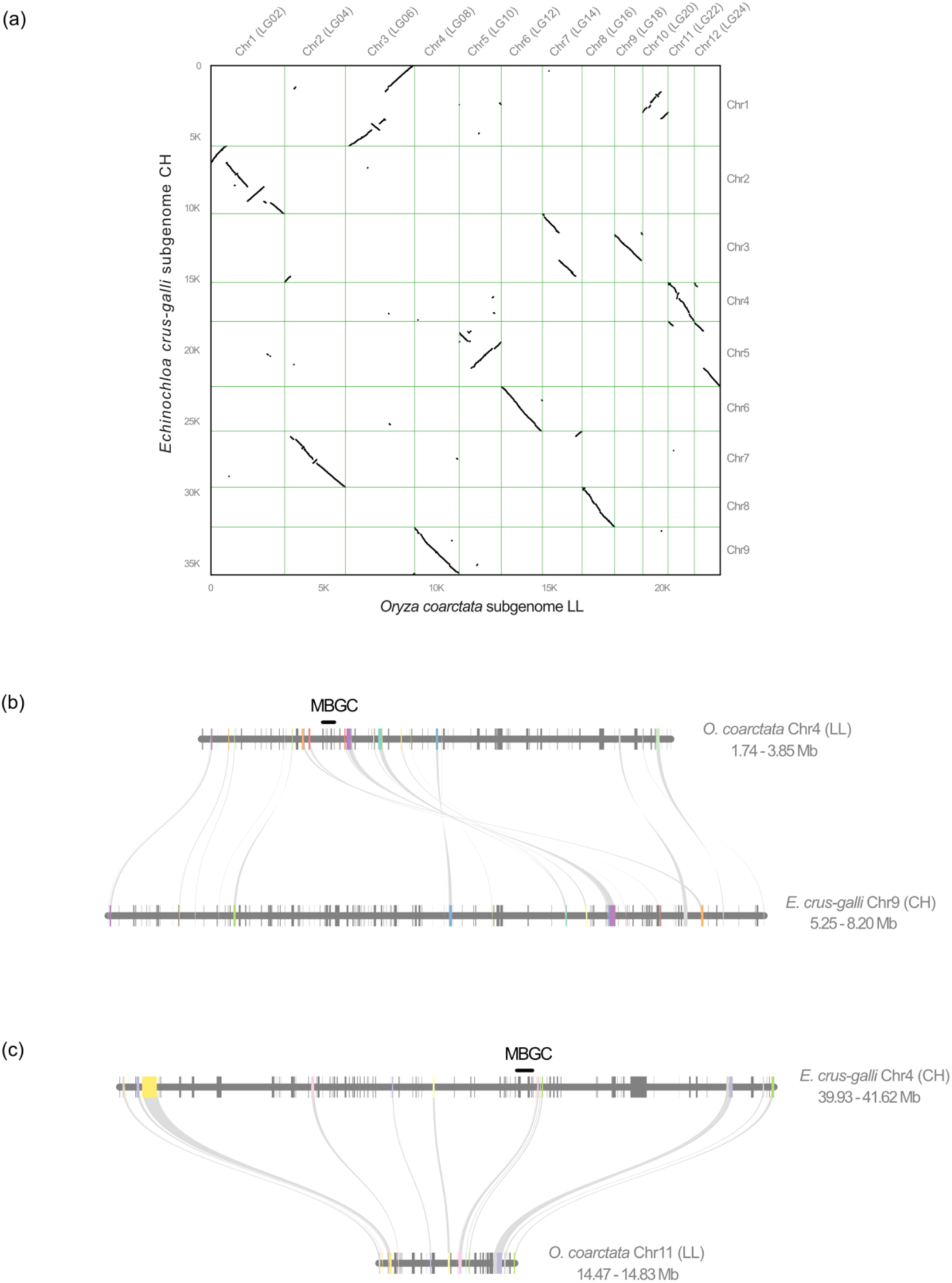
Synteny between *O. coarctata* sub-genome LL and *E. crus-galli* sub-genome CH showing the lack of synteny between the genomic region containing the MBGC in *O. coarctata* (Chr4) and *E. crus-galli* (Chr4 sub-genome CH). a) Dot plot showing whole-genome pairwise synteny between *O. coarctata* sub-genome LL and *E. crus-galli* sub-genome CH **b)** Synteny between the genomic region of *O. coarctata* containing MBGC (Chr4 LL) and its syntenic block in *E. crus-galli* sub-genome CH (Chr9). **c)** Synteny between the genomic region of *E. crus-galli* containing MBGC (Chr4 CH) and its syntenic block in *O. coarctata* sub-genome LL (Chr11). In (**b)** and **(c)**, grey lines connect the respective orthologs, and the black horizontal line indicates where the MBGC is located on each species.

**Supplementary Figure 7.**
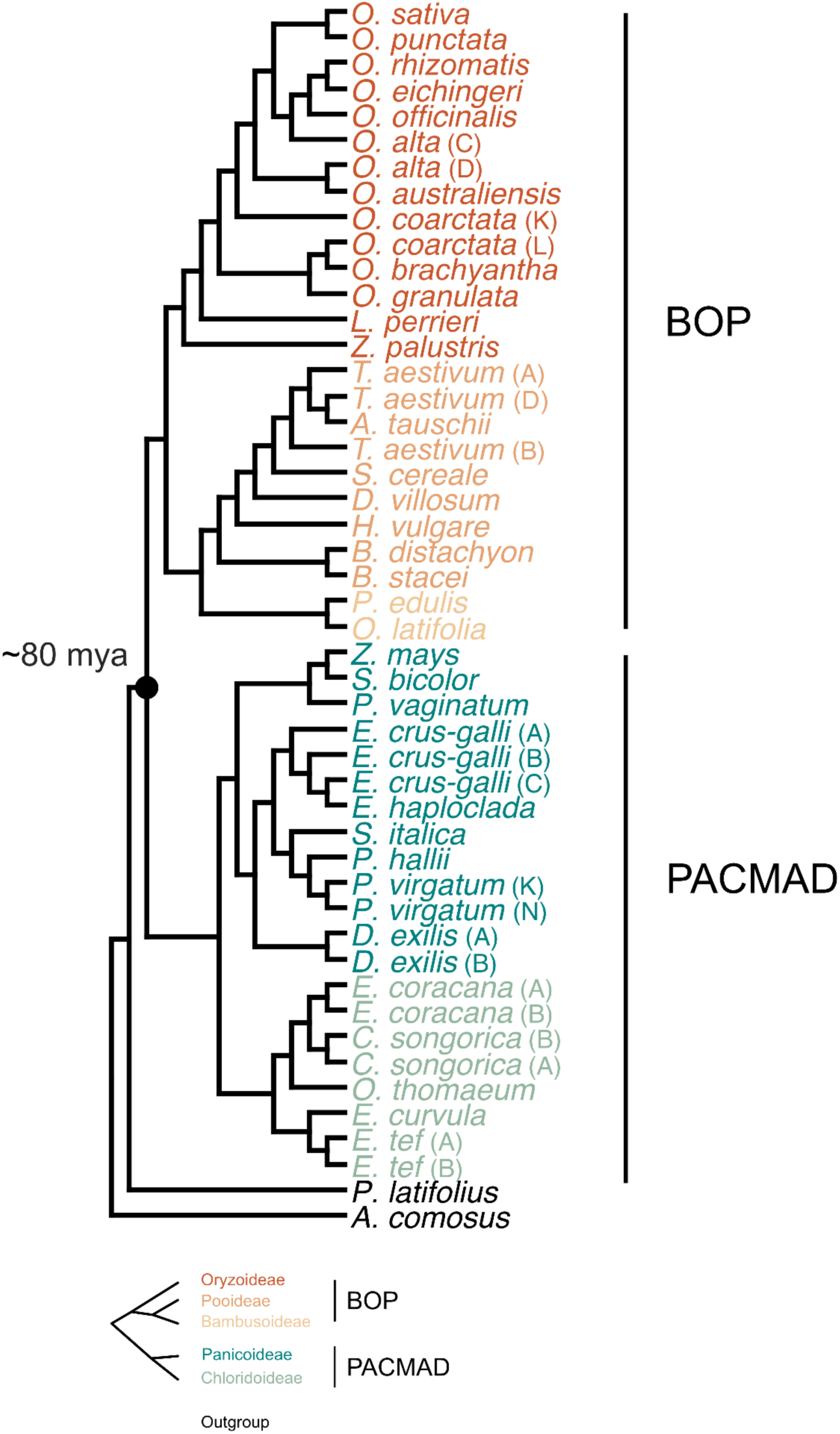
Phylogenetic relationship between the Poaceae species used in this study. The tree was inferred with the OrthoFinder algorithm STAG (Emms and Kelly, 2018) that uses all the species tree inferred from all orthogroups. *P. latifolius* (Poaceae) outgroup of core Poaceae (BOP and PACMAD), *A. comosus*, outgroup of Poaceae.

**Supplementary Figure 8.**
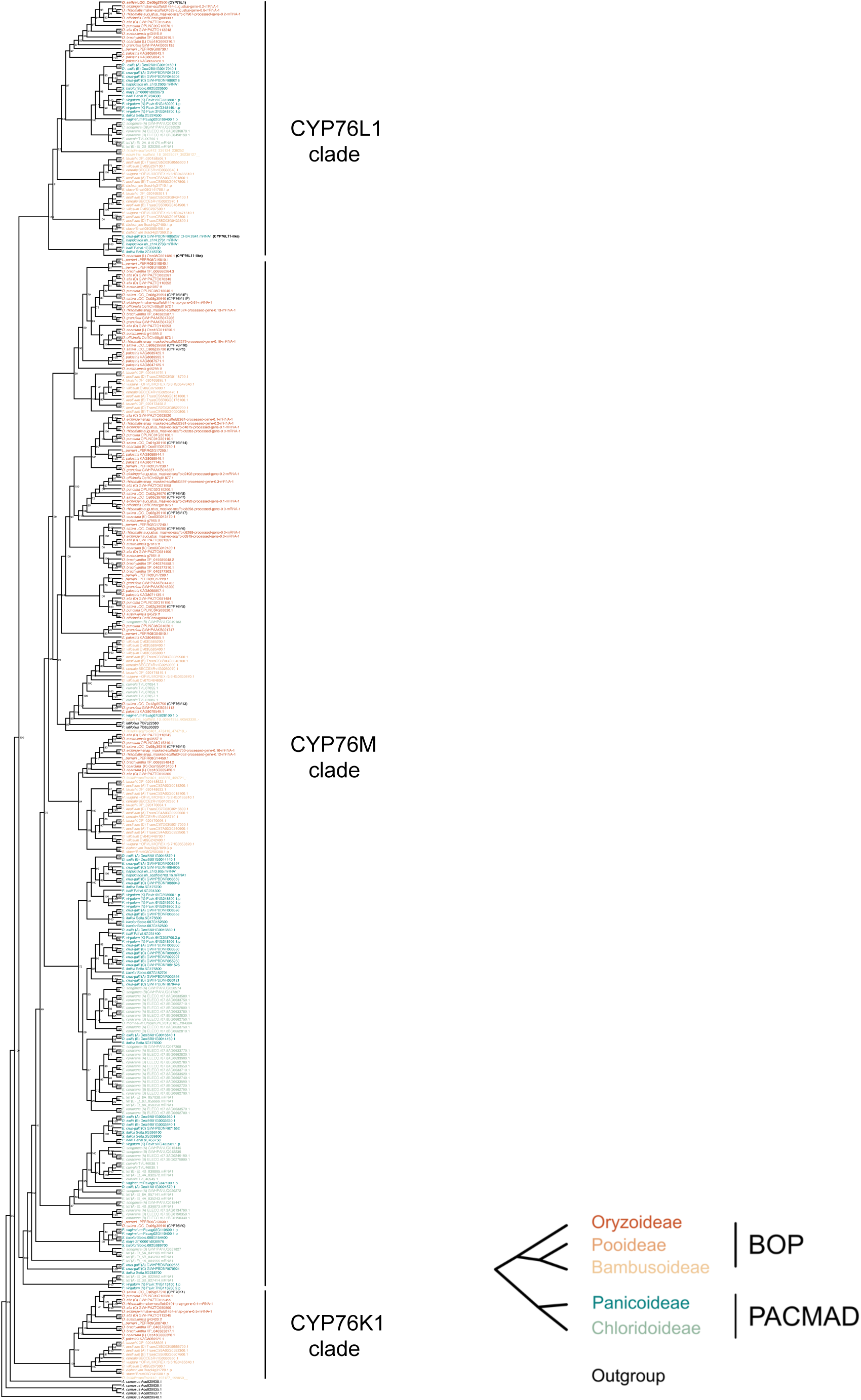
Cladogram representing the amino-acid-sequence maximum likelihood tree with the best-fit model selected by ModelFinder representing the relationship between the orthogroup containing *O. coarctat*a CYP76L11. The claudogram contains the CYP76K, CYP76L and CYP76M clades of 37 Poaceae species belonging to the subfamilies Oryzoideae, Pooideae, Bambusoideae, Panicoideae and Chloridoideae. The names of the different rice CYPs were taken from https://funricegenes.github.io/cytochrome_P450_monooxygenase_superfamily/. The phylogenetic relation between these subfamilies is represented in the bottom-right graph.

**Supplementary Figure 9.**
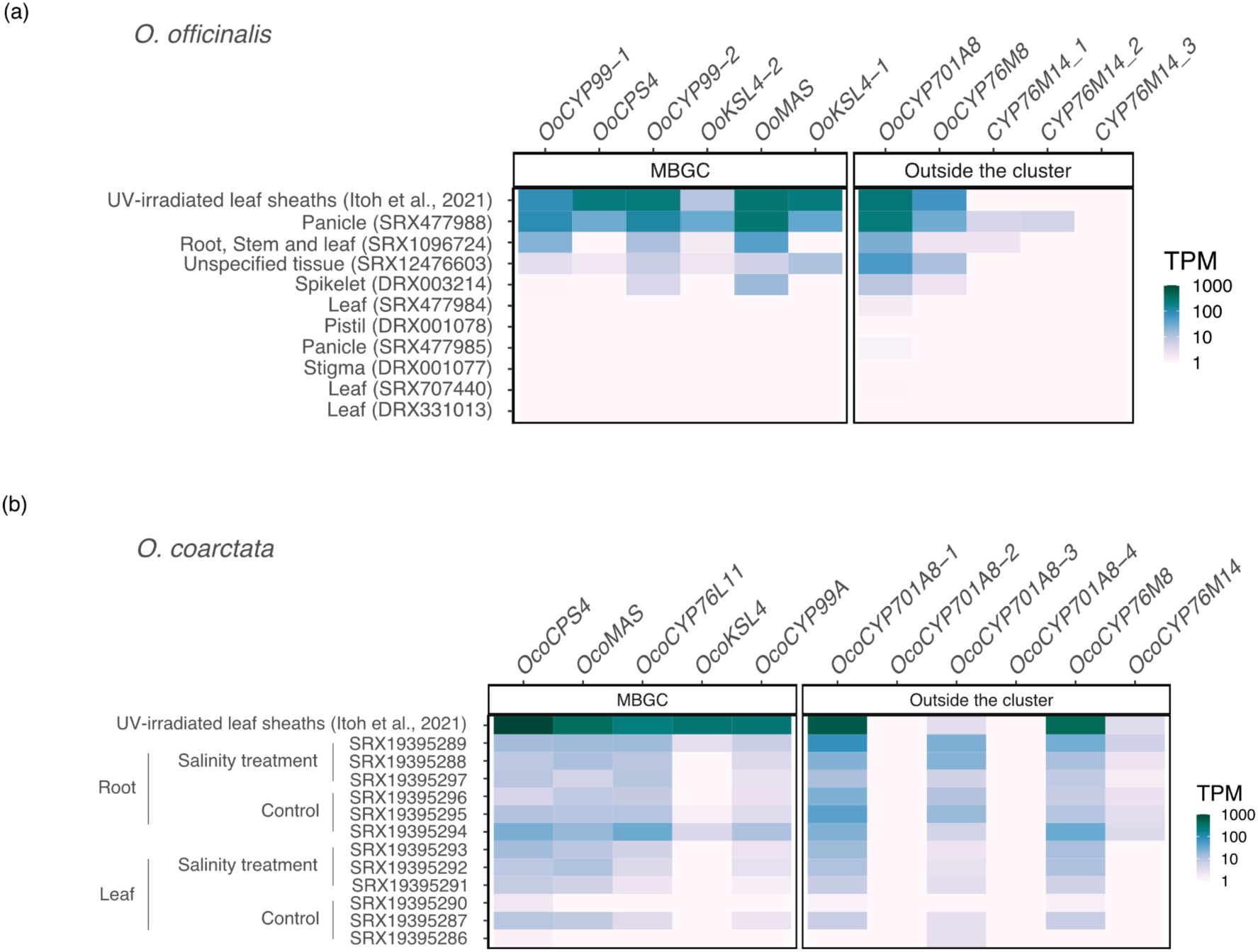
Expression of the momilactone biosynthetic orthologs within and outside the MBGC in a) *O. officinalis* and b) *O. coarctata*. The expression is represented as transcripts per million (TPM) and the colour is in logarithmic scale. The RNA-seq datasets were publicly available RNA-seq (see y axis on each heatmap) and these cover different tissues and treatments. Notice that the genes belonging to the MBGC are represented in the same order as in the cluster.

**Supplementary Figure 10.**
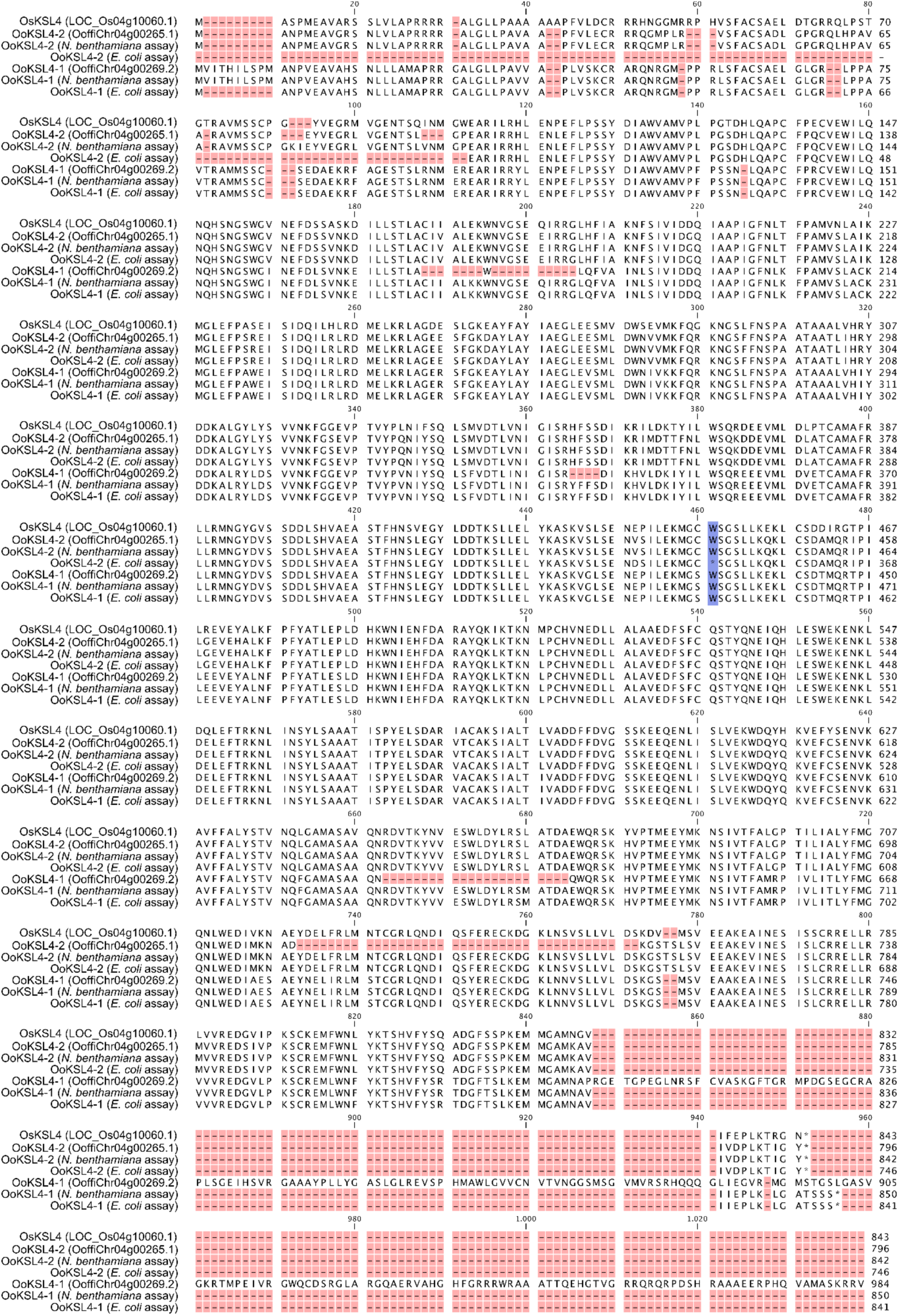
Amino acid sequence alignment of KSL4 orthologs. The alignment includes KSL4 from *O. sativa* (OsKSL4), its orthologs in *O. officinalis* from the annotation (KSL4-2: OoffiChr04g00265.1; KSL4-1: OoffiChr04g00269.2) and the cloned sequences used in *N. benthamiana* and *E. coli* assays. The position of the W445 clonal *de novo* mutation is highlighted in OoKSL4-2 (position assigned using the full length *OoKSL4-2* sequence cloned for the *N. benthamiana* transient expression as the reference sequence).

**Supplementary Figure 11.**
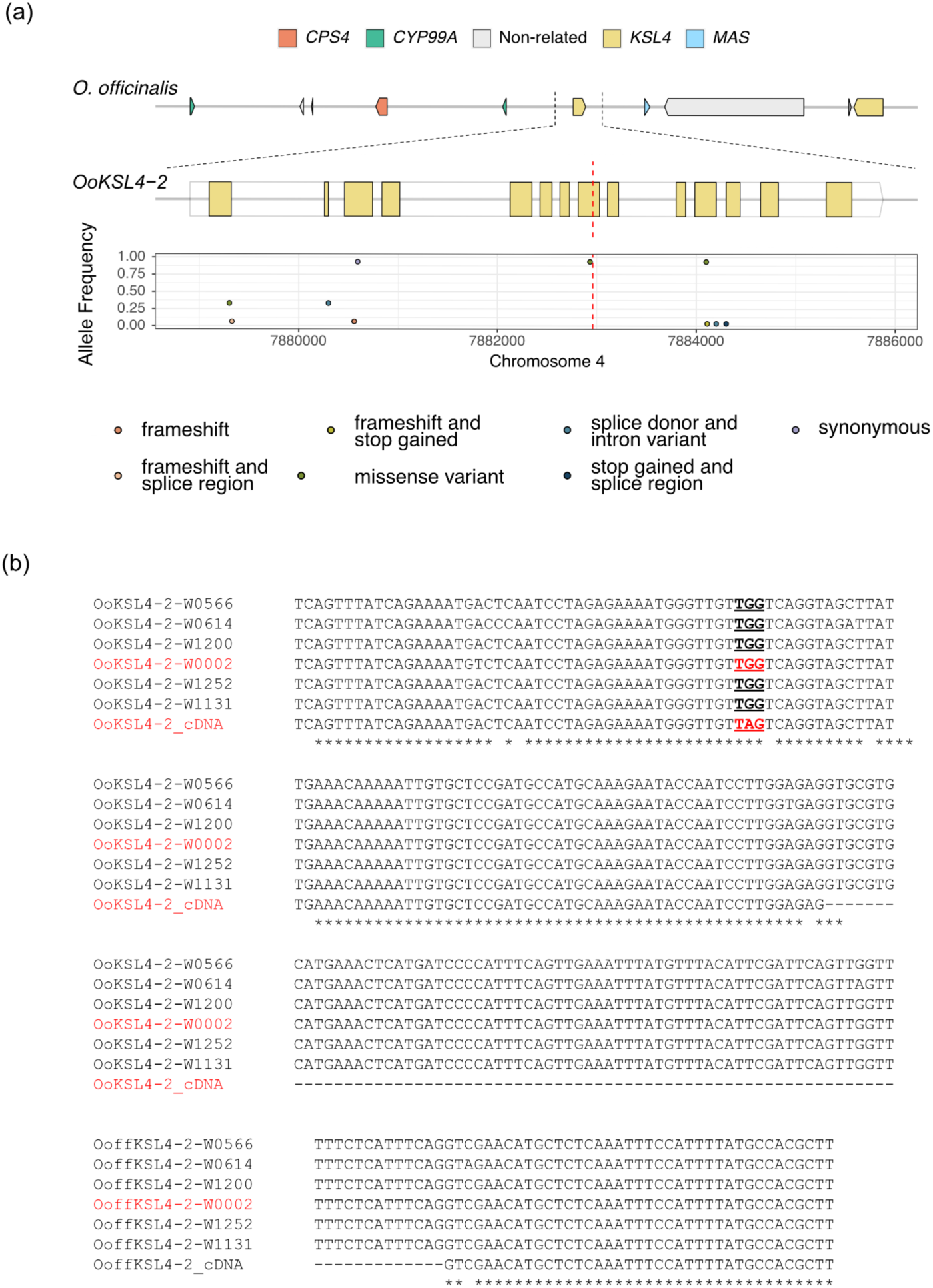
Prevalence of the nonsense G-to-A polymorphism in *OoKSL4-2* cDNA clones at the population level. **a)** The biosynthetic gene cluster (MBGC) from *O. officinalis*. Middle panel shows a magnification of the *OoKSL4-2* locus where the yellow filled rectangles represent exons of *OoKSL4-2*. Bottom panel shows the allele frequency of single nucleotide polymorphisms (SNPs) with a potential disruptive effect on *OoKSL4-2* from 15 *O. officinalis* accessions. The SNPs represented have allele frequency > 0.2 or a putative impact annotated as ‘high’. The red dashed line indicates the position of the G-to-A transition in the original cDNA clone, which is absent at the population level. **b)** Comparison of genome sequences of several *O. officinalis* accessions. The underlined codon indicates the mutation detected in the cDNA clones of OoKSL4-2 (W0002), which is absent in other accessions and in a different W0002 batch.

**Supplementary Figure 12.**
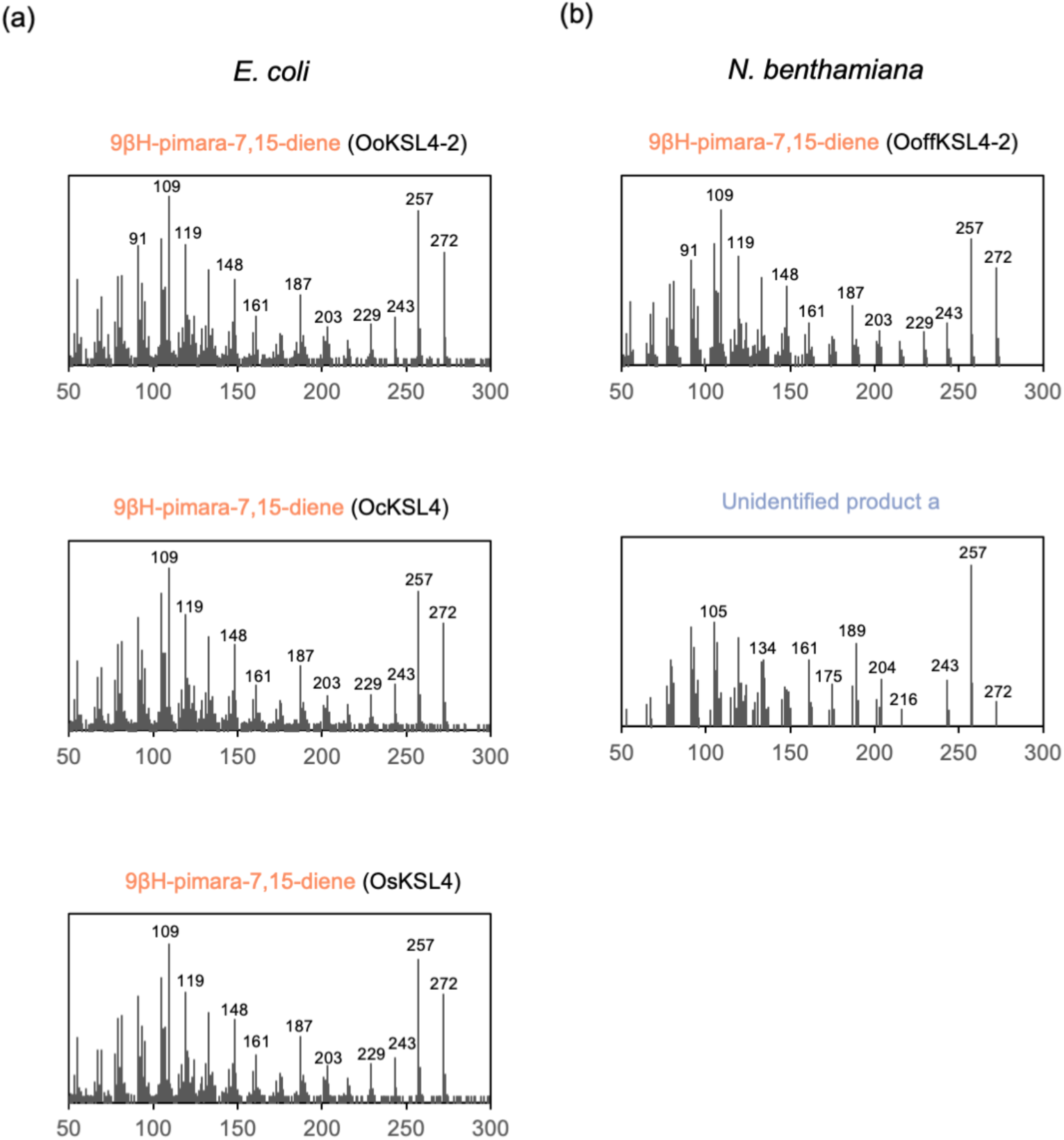
**Mass spectra of peaks identified in** *E. coli* metabolic engineering system (**a)** and *Nicotiana benthamiana* transient expression system **(b)**.

**Supplementary Figure 13.**
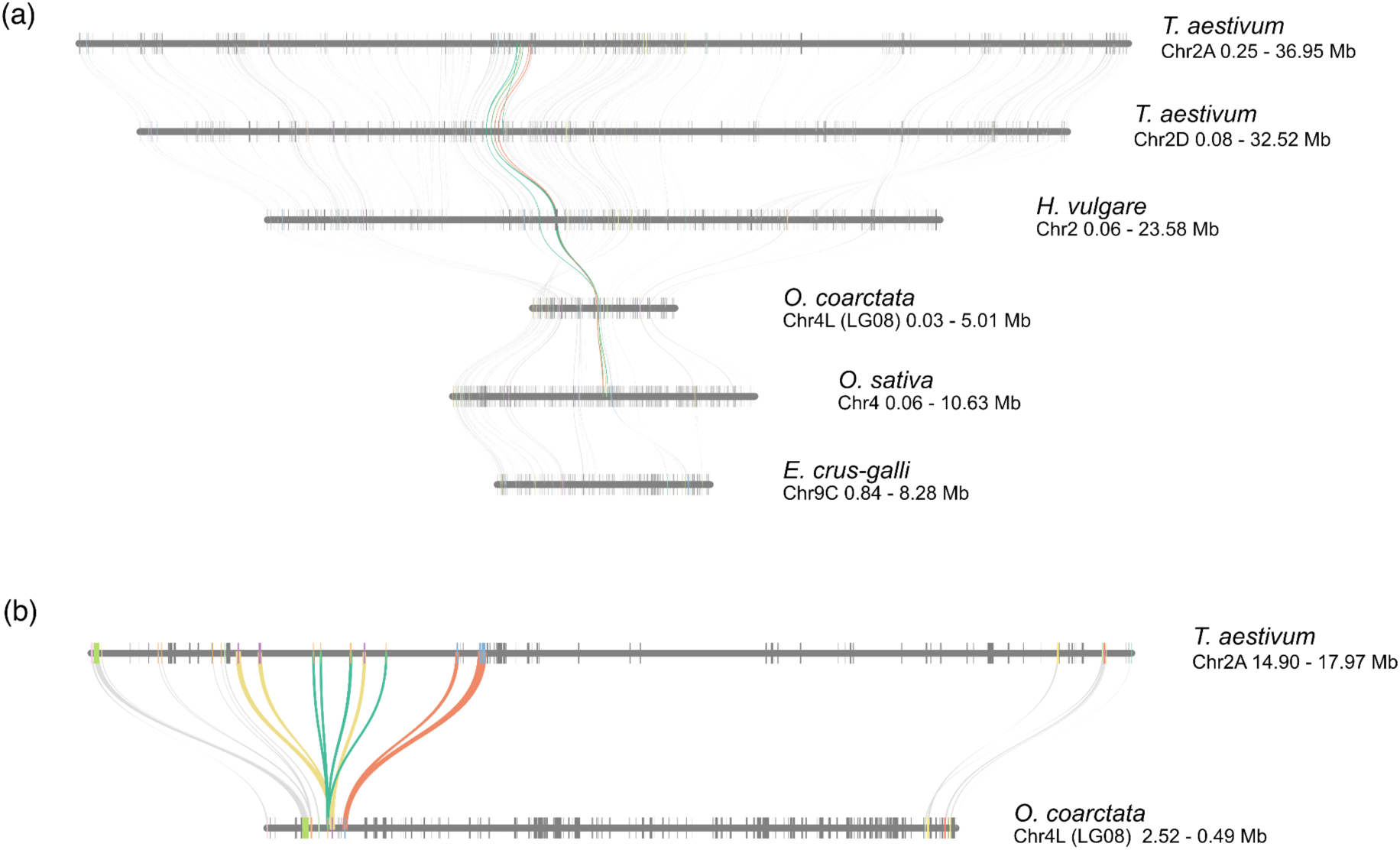
**Microsynteny analysis of the genomic regions containing the MBGC-like** from **a)** *Triticum aestivum* (wheat), *Hordeum vulgare* and the MBGC from *O. coarctata* and *O. sativa*. **b)** Magnification of the genomic region containing the MBGC-like and the MBGC in *T. aestivum* (sub-genome A) and *O. coarctata* showing the synteny between the flanking regions of both BGCs. Coloured lines represent MBGC-like orthologs and grey lines connect the respective orthologs.

### UPPLEMENTARY TABLES

**Supplementary Table 1.** Key statistics on the reference genome assemblies used in this study.

| Species | Version | Genome type | Assembly Size (Mb) | Annotated Loci | Scaffold N50 (Mb) | Complete BUSCOs (%) | Reference |
| --- | --- | --- | --- | --- | --- | --- | --- |
| *O. sativa* | v7.0 | AA | 375 | 42189 | 30 | 95 | Phytozome |
| *O. punctata* | v1.2 | BB | 394 | 31762 | 13 | 97,2 | Stein et al., 2018 |
| *O. officinalis* | GCA_008326285.1 | CC | 584 | 29930 | 0,51 | 94,6 or 98 | Shenton et al., 2020 |
| *O. rhizomatis* |  | CC | 559 | 32083 | 0,082 | 90,8 | Shenton et al., 2020 |
| *O. eichingeri* |  | CC | 471 | 3103 | 0,064 | 91,6 | Shenton et al., 2020 |
| *O. alta* | GWHAZTO000000000 | CCDD | 895 | 99312 | 37 | 98,2 | Yu et al., 2021 |
|  |  | sub-genome CC | 441 | 52861 |  |  |  |
|  |  | sub-genome DD | 435 | 46388 |  |  |  |
| *O. australiensis* | GCA_019925245.2 (MQ-UA-ANU_Oaus-KR_2.0) - Contig annotation | EE | 996 | 51057 | 1,9 | 91,9 | Phillips et al., 2022 |
| *O. coarctata* |  |  | 556 | 45571 | 23,1 | 96,22 | Zhao et al., 2023 |
|  |  | KK | 271 |  |  |  |  |
|  |  | LL | 261 |  |  |  |  |
| *O. brachyantha* | GCF_000231095.2 (ObraRS2) | FF | 261 | 32038 | 1,6 | 95,9 | Chen et al., 2013 |
| *O. granulata* | GWHAAKB000000000 | GG | 737 | 40131 | 0,92 | 96,53 | Shi et al., 2020 |
| *Leersia perrieri* | v1.4 | outgroup | 267 |  | 8,7 | 97,6 | Stein et al., 2018 |

**Supplementary Table 2.**
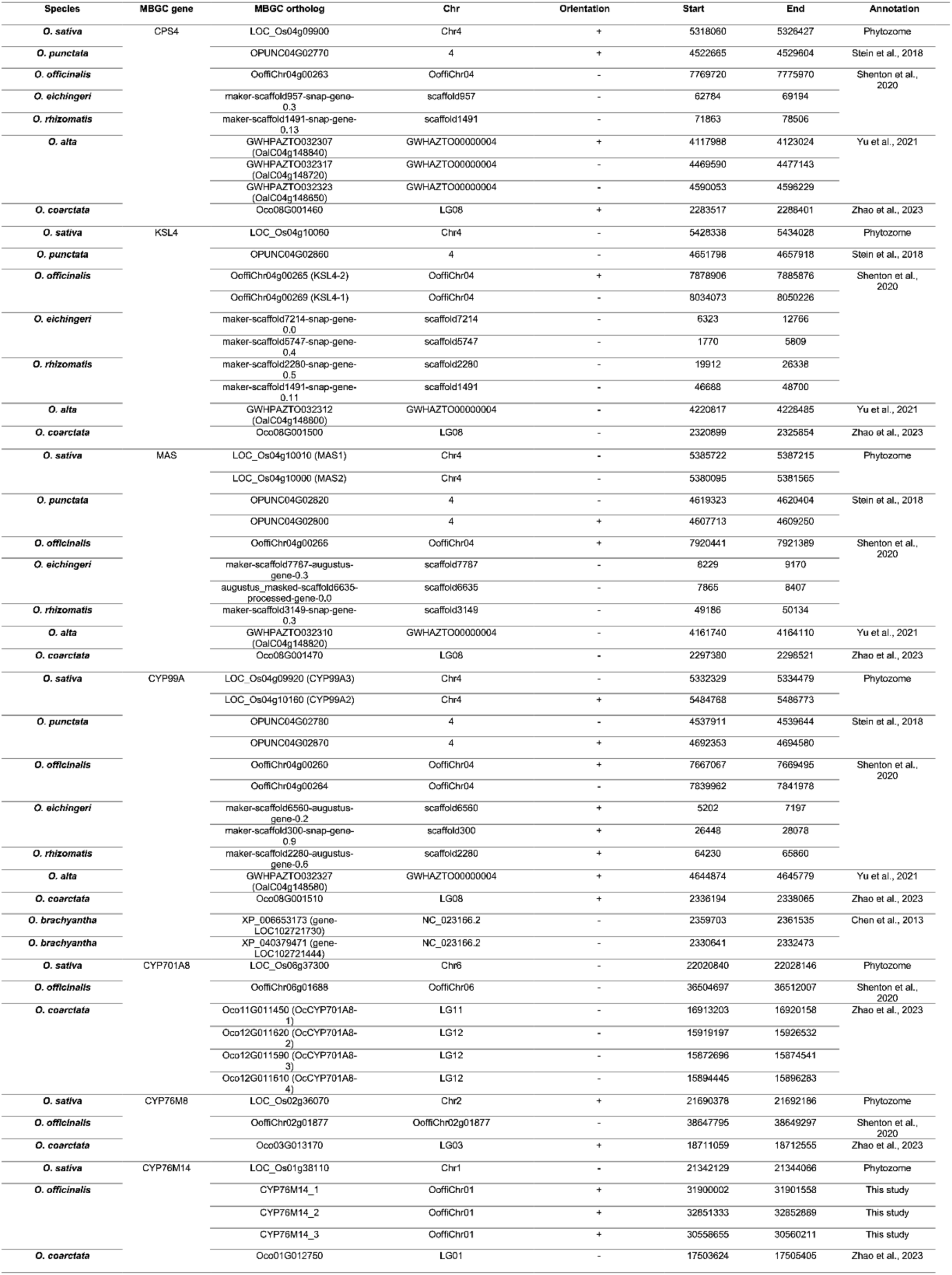
Scaffold and positional information on the orthologs of MBGC genes shown in Figure 1.

## Notes

### Competing Interest Statement

The authors have declared no competing interest.

### Summary of Updates

New momilactone measurement data for O. coarctata and O. officinalis; additional transcriptome data for the genes of the momilactone biosynthetic gene cluster from these two species; improved in vitro and N. benthamiana assays for KSL4 activity

## REFERENCES

Abrouk, M., Ahmed, H.I., Cubry, P., Šimoníková, D., Cauet, S., Pailles, Y., Bettgenhaeuser, J., Gapa, L., Scarcelli, N., Couderc, M., Zekraoui, L., Kathiresan, N., Čížková, J., Hřibová, E., Doležel, J., Arribat, S., Bergès, H., Wieringa, J.J., Gueye, M., Kane, N.A., Leclerc, C., Causse, S., Vancoppenolle, S., Billot, C., Wicker, T., Vigouroux, Y., Barnaud, A., Krattinger, S.G., 2020. Fonio millet genome unlocks African orphan crop diversity for agriculture in a changing climate. Nat. Commun. 11, 4488. 10.1038/s41467-020-18329-4

Ahuja, I., Kissen, R., Bones, A.M., 2012. Phytoalexins in defense against pathogens. Trends Plant Sci. 17, 73–90. 10.1016/j.tplants.2011.11.002

Bao, Y., Ge, S., 2004. Origin and phylogeny of Oryza species with the CD genome based on multiple-gene sequence data. Plant Syst. Evol. 249, 55–66. 10.1007/s00606-004-0173-8

Bennetzen, J.L., Schmutz, J., Wang, H., Percifield, R., Hawkins, J., Pontaroli, A.C., Estep, M., Feng, L., Vaughn, J.N., Grimwood, J., Jenkins, J., Barry, K., Lindquist, E., Hellsten, U., Deshpande, S., Wang, X., Wu, X., Mitros, T., Triplett, J., Yang, X., Ye, C.-Y., Mauro-Herrera, M., Wang, L., Li, P., Sharma, M., Sharma, R., Ronald, P.C., Panaud, O., Kellogg, E.A., Brutnell, T.P., Doust, A.N., Tuskan, G.A., Rokhsar, D., Devos, K.M., 2012. Reference genome sequence of the model plant Setaria. Nat. Biotechnol. 30, 555–561. 10.1038/nbt.2196

Cannarozzi, G., Plaza-Wüthrich, S., Esfeld, K., Larti, S., Wilson, Y.S., Girma, D., de Castro, E., Chanyalew, S., Blösch, R., Farinelli, L., Lyons, E., Schneider, M., Falquet, L., Kuhlemeier, C., Assefa, K., Tadele, Z., 2014. Genome and transcriptome sequencing identifies breeding targets in the orphan crop tef (Eragrostis tef). BMC Genomics 15, 581. 10.1186/1471-2164-15-581

Carballo, J., Santos, B. a. C.M., Zappacosta, D., Garbus, I., Selva, J.P., Gallo, C.A., Díaz, A., Albertini, E., Caccamo, M., Echenique, V., 2019. A high-quality genome of Eragrostis curvula grass provides insights into Poaceae evolution and supports new strategies to enhance forage quality. Sci. Rep. 9, 10250. 10.1038/s41598-019-46610-0

Chen, J., Huang, Q., Gao, D., Wang, Junyi, Lang, Y., Liu, T., Li, B., Bai, Z., Luis Goicoechea, J., Liang, C., Chen, C., Zhang, W., Sun, S., Liao, Y., Zhang, X., Yang, L., Song, C., Wang, M., Shi, J., Liu, G., Liu, J., Zhou, H., Zhou, W., Yu, Q., An, N., Chen, Y., Cai, Q., Wang, B., Liu, B., Min, J., Huang, Y., Wu, H., Li, Z., Zhang, Y., Yin, Y., Song, W., Jiang, J., Jackson, S.A., Wing, R.A., Wang, Jun, Chen, M., 2013. Whole-genome sequencing of Oryza brachyantha reveals mechanisms underlying Oryza genome evolution. Nat. Commun. 4, 1595. 10.1038/ncomms2596

Chen, S., Zhou, Y., Chen, Y., Gu, J., 2018. fastp: an ultra-fast all-in-one FASTQ preprocessor. Bioinformatics 34, i884–i890. 10.1093/bioinformatics/bty560

Cingolani, P., Platts, A., Wang, L.L., Coon, M., Nguyen, T., Wang, L., Land, S.J., Lu, X., Ruden, D.M., 2012. A program for annotating and predicting the effects of single nucleotide polymorphisms, SnpEff. Fly (Austin) 6, 80–92. 10.4161/fly.19695

Cock, P.J.A., Antao, T., Chang, J.T., Chapman, B.A., Cox, C.J., Dalke, A., Friedberg, I., Hamelryck, T., Kauff, F., Wilczynski, B., De Hoon, M.J.L., 2009. Biopython: freely available Python tools for computational molecular biology and bioinformatics. Bioinformatics 25, 1422–1423. 10.1093/bioinformatics/btp163

Danecek, P., Bonfield, J.K., Liddle, J., Marshall, J., Ohan, V., Pollard, M.O., Whitwham, A., Keane, T., McCarthy, S.A., Davies, R.M., Li, H., 2021. Twelve years of SAMtools and BCFtools. GigaScience 10, giab008. 10.1093/gigascience/giab008

De La Peña, R., Sattely, E.S., 2021. Rerouting plant terpene biosynthesis enables momilactone pathway elucidation. Nat. Chem. Biol. 17, 205–212. 10.1038/s41589-020-00669-3

Devos, K.M., Qi, P., Bahri, B.A., Gimode, D.M., Jenike, K., Manthi, S.J., Lule, D., Lux, T., Martinez-Bello, L., Pendergast, T.H., Plott, C., Saha, D., Sidhu, G.S., Sreedasyam, A., Wang, X., Wang, H., Wright, H., Zhao, J., Deshpande, S., de Villiers, S., Dida, M.M., Grimwood, J., Jenkins, J., Lovell, J., Mayer, K.F.X., Mneney, E.E., Ojulong, H.F., Schatz, M.C., Schmutz, J., Song, B., Tesfaye, K., Odeny, D.A., 2023. Genome analyses reveal population structure and a purple stigma color gene candidate in finger millet. Nat. Commun. 14, 3694. 10.1038/s41467-023-38915-6

Dobin, A., Davis, C.A., Schlesinger, F., Drenkow, J., Zaleski, C., Jha, S., Batut, P., Chaisson, M., Gingeras, T.R., 2013. STAR: ultrafast universal RNA-seq aligner. Bioinformatics 29, 15–21. 10.1093/bioinformatics/bts635

Dunning, L.T., Olofsson, J.K., Parisod, C., Choudhury, R.R., Moreno-Villena, J.J., Yang, Y., Dionora, J., Quick, W.P., Park, M., Bennetzen, J.L., Besnard, G., Nosil, P., Osborne, C.P., Christin, P.-A., 2019. Lateral transfers of large DNA fragments spread functional genes among grasses. Proc. Natl. Acad. Sci. 116, 4416–4425. 10.1073/pnas.1810031116

Emms, D.M., Kelly, S., 2019. OrthoFinder: phylogenetic orthology inference for comparative genomics. Genome Biol. 20, 238. 10.1186/s13059-019-1832-y

Emms, D.M., Kelly, S., 2018. STAG: Species Tree Inference from All Genes. 10.1101/267914

Emms, D.M., Kelly, S., 2015. OrthoFinder: solving fundamental biases in whole genome comparisons dramatically improves orthogroup inference accuracy. Genome Biol. 16, 157. 10.1186/s13059-015-0721-2

Ewels, P.A., Peltzer, A., Fillinger, S., Patel, H., Alneberg, J., Wilm, A., Garcia, M.U., Di Tommaso, P., Nahnsen, S., 2020. The nf-core framework for community-curated bioinformatics pipelines. Nat. Biotechnol. 38, 276–278. 10.1038/s41587-020-0439-x

Frerebeau, N., 2023. khroma: Colour Schemes for Scientific Data Visualization. Zenodo. 10.5281/zenodo.8269306

Garcia, M., Juhos, S., Larsson, M., Olason, P.I., Martin, M., Eisfeldt, J., DiLorenzo, S., Sandgren, J., Ståhl, T.D.D., Ewels, P., Wirta, V., Nistér, M., Käller, M., Nystedt, B., 2020. Sarek: A portable workflow for whole-genome sequencing analysis of germline and somatic variants. F1000Research. 10.12688/f1000research.16665.2

Ge, S., Sang, T., Lu, B.-R., Hong, D.-Y., 1999. Phylogeny of rice genomes with emphasis on origins of allotetraploid species. Proc. Natl. Acad. Sci. U. S. A. 96, 14400–14405.

Gordon, S.P., Contreras-Moreira, B., Levy, J.J., Djamei, A., Czedik-Eysenberg, A., Tartaglio, V.S., Session, A., Martin, J., Cartwright, A., Katz, A., Singan, V.R., Goltsman, E., Barry, K., Dinh-Thi, V.H., Chalhoub, B., Diaz-Perez, A., Sancho, R., Lusinska, J., Wolny, E., Nibau, C., Doonan, J.H., Mur, L.A.J., Plott, C., Jenkins, J., Hazen, S.P., Lee, S.J., Shu, S., Goodstein, D., Rokhsar, D., Schmutz, J., Hasterok, R., Catalan, P., Vogel, J.P., 2020. Gradual polyploid genome evolution revealed by pan-genomic analysis of Brachypodium hybridum and its diploid progenitors. Nat. Commun. 11, 3670. 10.1038/s41467-020-17302-5

Guo, L., Qiu, J., Ye, C., Jin, G., Mao, L., Zhang, H., Yang, X., Peng, Q., Wang, Yingying, Jia, L., Lin, Z., Li, G., Fu, F., Liu, C., Chen, L., Shen, E., Wang, W., Chu, Q., Wu, D., Wu, S., Xia, C., Zhang, Y., Zhou, X., Wang, L., Wu, L., Song, W., Wang, Yunfei, Shu, Q., Aoki, D., Yumoto, E., Yokota, T., Miyamoto, K., Okada, K., Kim, D.-S., Cai, D., Zhang, C., Lou, Y., Qian, Q., Yamaguchi, H., Yamane, H., Kong, C.-H., Timko, M.P., Bai, L., Fan, L., 2017. Echinochloa crus-galli genome analysis provides insight into its adaptation and invasiveness as a weed. Nat. Commun. 8, 1031. 10.1038/s41467-017-01067-5

Guo, Y.-L., Ge, S., 2005. Molecular phylogeny of Oryzeae (Poaceae) based on DNA sequences from chloroplast, mitochondrial, and nuclear genomes. Am. J. Bot. 92, 1548–1558. 10.3732/ajb.92.9.1548

Guo, Z.-H., Ma, P.-F., Yang, G.-Q., Hu, J.-Y., Liu, Y.-L., Xia, E.-H., Zhong, M.-C., Zhao, L., Sun, G.-L., Xu, Y.-X., Zhao, Y.-J., Zhang, Y.-C., Zhang, Y.-X., Zhang, X.-M., Zhou, M.-Y., Guo, Y., Guo, C., Liu, J.-X., Ye, X.-Y., Chen, Y.-M., Yang, Y., Han, B., Lin, C.-S., Lu, Y., Li, D.-Z., 2019. Genome Sequences Provide Insights into the Reticulate Origin and Unique Traits of Woody Bamboos. Mol. Plant 12, 1353–1365. 10.1016/j.molp.2019.05.009

Haas, M., Kono, T., Macchietto, M., Millas, R., McGilp, L., Shao, M., Duquette, J., Qiu, Y., Hirsch, C.N., Kimball, J., 2021. Whole-genome assembly and annotation of northern wild rice, Zizania palustris L., supports a whole-genome duplication in the Zizania genus. Plant J. 107, 1802–1818. 10.1111/tpj.15419

Hibdige, S.G.S., Raimondeau, P., Christin, P.-A., Dunning, L.T., 2021. Widespread lateral gene transfer among grasses. New Phytol. 230, 2474–2486. 10.1111/nph.17328

Hoang, D.T., Chernomor, O., von Haeseler, A., Minh, B.Q., Vinh, L.S., 2018. UFBoot2: Improving the Ultrafast Bootstrap Approximation. Mol. Biol. Evol. 35, 518–522. 10.1093/molbev/msx281

Itoh, A., Nakazato, S., Wakabayashi, H., Hamano, A., Shenton, M.R., Miyamoto, K., Mitsuhashi, W., Okada, K., Toyomasu, T., 2021. Functional kaurene-synthase-like diterpene synthases lacking a gamma domain are widely present in Oryza and related species. Biosci. Biotechnol. Biochem. 85, 1945–1952. 10.1093/bbb/zbab127

Kajiya-Kanegae, H., Ohyanagi, H., Ebata, T., Tanizawa, Y., Onogi, A., Sawada, Y., Hirai, M.Y., Wang, Z.-X., Han, B., Toyoda, A., Fujiyama, A., Iwata, H., Tsuda, K., Suzuki, T., Nosaka-Takahashi, M., Nonomura, K., Nakamura, Y., Kawamoto, S., Kurata, N., Sato, Y., 2021. OryzaGenome2.1: Database of Diverse Genotypes in Wild Oryza Species. Rice 14, 24. 10.1186/s12284-021-00468-x

Kalyaanamoorthy, S., Minh, B.Q., Wong, T.K.F., von Haeseler, A., Jermiin, L.S., 2017. ModelFinder: fast model selection for accurate phylogenetic estimates. Nat. Methods 14, 587–589. 10.1038/nmeth.4285

Kato, T., Kabuto, C., Sasaki, N., Tsunagawa, M., Aizawa, H., Fujita, K., Kato, Y., Kitahara, Y., Takahashi, N., 1973. Momilactones, growth inhibitors from rice, oryza sativa L. Tetrahedron Lett. 14, 3861–3864. 10.1016/S0040-4039(01)87058-1

Katoh, K., Misawa, K., Kuma, K., Miyata, T., 2002. MAFFT: a novel method for rapid multiple sequence alignment based on fast Fourier transform. Nucleic Acids Res. 30, 3059–3066. 10.1093/nar/gkf436

Katoh, K., Standley, D.M., 2013. MAFFT Multiple Sequence Alignment Software Version 7: Improvements in Performance and Usability. Mol. Biol. Evol. 30, 772–780. 10.1093/molbev/mst010

Kato-Noguchi, H., Hasegawa, M., Ino, T., Ota, K., Kujime, H., 2010. Contribution of momilactone A and B to rice allelopathy. J. Plant Physiol. 167, 787–791. 10.1016/j.jplph.2010.01.014

Kato-Noguchi, H., Ino, T., 2003. Rice seedlings release momilactone B into the environment. Phytochemistry 63, 551–554. 10.1016/S0031-9422(03)00194-8

Kitaoka, N., Zhang, J., Oyagbenro, R.K., Brown, B., Wu, Y., Yang, B., Li, Z., Peters, R.J., 2021. Interdependent evolution of biosynthetic gene clusters for momilactone production in rice. Plant Cell 33, 290–305. 10.1093/plcell/koaa023

Kodama, O., Suzuki, T., Miyakawa, J., Akatsuka, T., 1988. Ultraviolet-Induced Accumulation of Phytoalexins in Rice Leaves. Agric. Biol. Chem. 52, 2469–2473. 10.1080/00021369.1988.10869067

Kraehmer, H., Jabran, K., Mennan, H., Chauhan, B.S., 2016. Global distribution of rice weeds – A review. Crop Prot. 80, 73–86. 10.1016/j.cropro.2015.10.027

Krueger, F., James, F., Ewels, P., Afyounian, E., Schuster-Boeckler, B., 2021. FelixKrueger/TrimGalore: v0.6.7 - DOI via Zenodo. 10.5281/zenodo.5127899

Liu, Y., Balcke, G.U., Porzel, A., Mahdi, L., Scherr-Henning, A., Bathe, U., Zuccaro, A., Tissier, A., 2021. A barley gene cluster for the biosynthesis of diterpenoid phytoalexins. bioRxiv. 10.1101/2021.05.21.445084

Liu, Z., Cheema, J., Vigouroux, M., Hill, L., Reed, J., Paajanen, P., Yant, L., Osbourn, A., 2020. Formation and diversification of a paradigm biosynthetic gene cluster in plants. Nat. Commun. 11, 5354. 10.1038/s41467-020-19153-6

Lovell, J.T., Jenkins, J., Lowry, D.B., Mamidi, S., Sreedasyam, A., Weng, X., Barry, K., Bonnette, J., Campitelli, B., Daum, C., Gordon, S.P., Gould, B.A., Khasanova, A., Lipzen, A., MacQueen, A., Palacio-Mejía, J.D., Plott, C., Shakirov, E.V., Shu, S., Yoshinaga, Y., Zane, M., Kudrna, D., Talag, J.D., Rokhsar, D., Grimwood, J., Schmutz, J., Juenger, T.E., 2018. The genomic landscape of molecular responses to natural drought stress in Panicum hallii. Nat. Commun. 9, 5213. 10.1038/s41467-018-07669-x

Lovell, J.T., MacQueen, A.H., Mamidi, S., Bonnette, J., Jenkins, J., Napier, J.D., Sreedasyam, A., Healey, A., Session, A., Shu, S., Barry, K., Bonos, S., Boston, L., Daum, C., Deshpande, S., Ewing, A., Grabowski, P.P., Haque, T., Harrison, M., Jiang, J., Kudrna, D., Lipzen, A., Pendergast, T.H., Plott, C., Qi, P., Saski, C.A., Shakirov, E.V., Sims, D., Sharma, M., Sharma, R., Stewart, A., Singan, V.R., Tang, Y., Thibivillier, S., Webber, J., Weng, X., Williams, M., Wu, G.A., Yoshinaga, Y., Zane, M., Zhang, L., Zhang, J., Behrman, K.D., Boe, A.R., Fay, P.A., Fritschi, F.B., Jastrow, J.D., Lloyd-Reilley, J., Martínez-Reyna, J.M., Matamala, R., Mitchell, R.B., Rouquette, F.M., Ronald, P., Saha, M., Tobias, C.M., Udvardi, M., Wing, R.A., Wu, Y., Bartley, L.E., Casler, M., Devos, K.M., Lowry, D.B., Rokhsar, D.S., Grimwood, J., Juenger, T.E., Schmutz, J., 2021. Genomic mechanisms of climate adaptation in polyploid bioenergy switchgrass. Nature 590, 438–444. 10.1038/s41586-020-03127-1

Lu, F., Ammiraju, J.S.S., Sanyal, A., Zhang, S., Song, R., Chen, J., Li, G., Sui, Y., Song, X., Cheng, Z., Oliveira, A.C. de, Bennetzen, J.L., Jackson, S.A., Wing, R.A., Chen, M., 2009. Comparative sequence analysis of MONOCULM1-orthologous regions in 14 Oryza genomes. Proc. Natl. Acad. Sci. 106, 2071–2076. 10.1073/pnas.0812798106

Ma, P.-F., Liu, Y.-L., Jin, G.-H., Liu, J.-X., Wu, H., He, J., Guo, Z.-H., Li, D.-Z., 2021. The Pharus latifolius genome bridges the gap of early grass evolution. Plant Cell 33, 846–864. 10.1093/plcell/koab015

Mascher, M., Wicker, T., Jenkins, J., Plott, C., Lux, T., Koh, C.S., Ens, J., Gundlach, H., Boston, L.B., Tulpová, Z., Holden, S., Hernández-Pinzón, I., Scholz, U., Mayer, K.F.X., Spannagl, M., Pozniak, C.J., Sharpe, A.G., Šimková, H., Moscou, M.J., Grimwood, J., Schmutz, J., Stein, N., 2021. Long-read sequence assembly: a technical evaluation in barley. Plant Cell 33, 1888–1906. 10.1093/plcell/koab077

McCormick, R.F., Truong, S.K., Sreedasyam, A., Jenkins, J., Shu, S., Sims, D., Kennedy, M., Amirebrahimi, M., Weers, B.D., McKinley, B., Mattison, A., Morishige, D.T., Grimwood, J., Schmutz, J., Mullet, J.E., 2018. The Sorghum bicolor reference genome: improved assembly, gene annotations, a transcriptome atlas, and signatures of genome organization. Plant J. 93, 338–354. 10.1111/tpj.13781

McKenna, A., Hanna, M., Banks, E., Sivachenko, A., Cibulskis, K., Kernytsky, A., Garimella, K., Altshuler, D., Gabriel, S., Daly, M., DePristo, M.A., 2010. The Genome Analysis Toolkit: A MapReduce framework for analyzing next-generation DNA sequencing data. Genome Res. 20, 1297–1303. 10.1101/gr.107524.110

Ming, R., VanBuren, R., Wai, C.M., Tang, H., Schatz, M.C., Bowers, J.E., Lyons, E., Wang, M.-L., Chen, J., Biggers, E., Zhang, Jisen, Huang, L., Zhang, L., Miao, W., Zhang, Jian, Ye, Z., Miao, C., Lin, Z., Wang, H., Zhou, H., Yim, W.C., Priest, H.D., Zheng, C., Woodhouse, M., Edger, P.P., Guyot, R., Guo, H.-B., Guo, H., Zheng, G., Singh, R., Sharma, A., Min, X., Zheng, Y., Lee, H., Gurtowski, J., Sedlazeck, F.J., Harkess, A., McKain, M.R., Liao, Z., Fang, J., Liu, J., Zhang, X., Zhang, Q., Hu, W., Qin, Y., Wang, K., Chen, L.-Y., Shirley, N., Lin, Y.-R., Liu, L.-Y., Hernandez, A.G., Wright, C.L., Bulone, V., Tuskan, G.A., Heath, K., Zee, F., Moore, P.H., Sunkar, R., Leebens-Mack, J.H., Mockler, T., Bennetzen, J.L., Freeling, M., Sankoff, D., Paterson, A.H., Zhu, X., Yang, X., Smith, J.A.C., Cushman, J.C., Paull, R.E., Yu, Q., 2015. The pineapple genome and the evolution of CAM photosynthesis. Nat. Genet. 47, 1435–1442. 10.1038/ng.3435

Minh, B.Q., Schmidt, H.A., Chernomor, O., Schrempf, D., Woodhams, M.D., von Haeseler, A., Lanfear, R., 2020. IQ-TREE 2: New Models and Efficient Methods for Phylogenetic Inference in the Genomic Era. Mol. Biol. Evol. 37, 1530–1534. 10.1093/molbev/msaa015

Miyamoto, K., Fujita, M., Shenton, M.R., Akashi, S., Sugawara, C., Sakai, A., Horie, K., Hasegawa, M., Kawaide, H., Mitsuhashi, W., Nojiri, H., Yamane, H., Kurata, N., Okada, K., Toyomasu, T., 2016. Evolutionary trajectory of phytoalexin biosynthetic gene clusters in rice. Plant J. 87, 293–304. 10.1111/tpj.13200

Nützmann, H.-W., Huang, A., Osbourn, A., 2016. Plant metabolic clusters – from genetics to genomics. New Phytol. 211, 771–789. 10.1111/nph.13981

Okada, K., 2011. The Biosynthesis of Isoprenoids and the Mechanisms Regulating It in Plants. Biosci. Biotechnol. Biochem. 75, 1219–1225. 10.1271/bbb.110228

Otomo, K., Kanno, Y., Motegi, A., Kenmoku, H., Yamane, H., Mitsuhashi, W., Oikawa, H., Toshima, H., Itoh, H., Matsuoka, M., Sassa, T., Toyomasu, T., 2004a. Diterpene Cyclases Responsible for the Biosynthesis of Phytoalexins, Momilactones A, B, and Oryzalexins A–F in Rice. Biosci. Biotechnol. Biochem. 68, 2001–2006. 10.1271/bbb.68.2001

Otomo, K., Kenmoku, H., Oikawa, H., König, W.A., Toshima, H., Mitsuhashi, W., Yamane, H., Sassa, T., Toyomasu, T., 2004b. Biological functions of ent- and syn-copalyl diphosphate synthases in rice: key enzymes for the branch point of gibberellin and phytoalexin biosynthesis. Plant J. 39, 886–893. 10.1111/j.1365-313X.2004.02175.x

Ouyang, S., Zhu, W., Hamilton, J., Lin, H., Campbell, M., Childs, K., Thibaud-Nissen, F., Malek, R.L., Lee, Y., Zheng, L., Orvis, J., Haas, B., Wortman, J., Buell, C.R., 2007. The TIGR Rice Genome Annotation Resource: improvements and new features. Nucleic Acids Res. 35, D883–D887. 10.1093/nar/gkl976

Patel, H., Ewels, P., Peltzer, A., Manning, J., Botvinnik, O., Sturm, G., Garcia, M.U., Moreno, D., Vemuri, P., bot, nf-core, Binzer-Panchal, M., silviamorins, Pantano, L., Zepper, M., Syme, R., Talbot, A., Kelly, G., Hanssen, F., Yates, J.A.F., Espinosa-Carrasco, J., rfenouil, Cheshire, C., marchoeppner, Miller, E., Zhou, P., Guinchard, S., Gabernet, G., Mertes, C., Straub, D., Tommaso, P.D., 2024. nf-core/rnaseq: nf-core/rnaseq v3.14.0 - Hassium Honey Badger. 10.5281/zenodo.10471647

Patro, R., Duggal, G., Love, M.I., Irizarry, R.A., Kingsford, C., 2017. Salmon provides fast and bias-aware quantification of transcript expression. Nat. Methods 14, 417–419. 10.1038/nmeth.4197

Pertea, G., Pertea, M., 2020. GFF Utilities: GffRead and GffCompare. F1000Research. 10.12688/f1000research.23297.1

Phillips, A.L., Ferguson, S., Watson-Haigh, N.S., Jones, A.W., Borevitz, J.O., Burton, R.A., Atwell, B.J., 2022. The first long-read nuclear genome assembly of Oryza australiensis, a wild rice from northern Australia. Sci. Rep. 12, 10823. 10.1038/s41598-022-14893-5

Polturak, G., Dippe, M., Stephenson, M.J., Chandra Misra, R., Owen, C., Ramirez-Gonzalez, R.H., Haidoulis, J.F., Schoonbeek, H.-J., Chartrain, L., Borrill, P., Nelson, D.R., Brown, J.K.M., Nicholson, P., Uauy, C., Osbourn, A., 2022a. Pathogen-induced biosynthetic pathways encode defense-related molecules in bread wheat. Proc. Natl. Acad. Sci. 119, e2123299119. 10.1073/pnas.2123299119

Polturak, G., Liu, Z., Osbourn, A., 2022b. New and emerging concepts in the evolution and function of plant biosynthetic gene clusters. Curr. Opin. Green Sustain. Chem. 33, 100568. 10.1016/j.cogsc.2021.100568

Poplin, R., Ruano-Rubio, V., DePristo, M.A., Fennell, T.J., Carneiro, M.O., Auwera, G.A.V. der, Kling, D.E., Gauthier, L.D., Levy-Moonshine, A., Roazen, D., Shakir, K., Thibault, J., Chandran, S., Whelan, C., Lek, M., Gabriel, S., Daly, M.J., Neale, B., MacArthur, D.G., Banks, E., 2018. Scaling accurate genetic variant discovery to tens of thousands of samples. bioRxiv 201178. 10.1101/201178

Rabanus-Wallace, M.T., Hackauf, B., Mascher, M., Lux, T., Wicker, T., Gundlach, H., Baez, M., Houben, A., Mayer, K.F.X., Guo, L., Poland, J., Pozniak, C.J., Walkowiak, S., Melonek, J., Praz, C.R., Schreiber, M., Budak, H., Heuberger, M., Steuernagel, B., Wulff, B., Börner, A., Byrns, B., Čížková, J., Fowler, D.B., Fritz, A., Himmelbach, A., Kaithakottil, G., Keilwagen, J., Keller, B., Konkin, D., Larsen, J., Li, Q., Myśków, B., Padmarasu, S., Rawat, N., Sesiz, U., Biyiklioglu-Kaya, S., Sharpe, A., Šimková, H., Small, I., Swarbreck, D., Toegelová, H., Tsvetkova, N., Voylokov, A.V., Vrána, J., Bauer, E., Bolibok-Bragoszewska, H., Doležel, J., Hall, A., Jia, J., Korzun, V., Laroche, A., Ma, X.-F., Ordon, F., Özkan, H., Rakoczy-Trojanowska, M., Scholz, U., Schulman, A.H., Siekmann, D., Stojałowski, S., Tiwari, V.K., Spannagl, M., Stein, N., 2021. Chromosome-scale genome assembly provides insights into rye biology, evolution and agronomic potential. Nat. Genet. 53, 564–573. 10.1038/s41588-021-00807-0

Schnable, P.S., Ware, D., Fulton, R.S., Stein, J.C., Wei, F., Pasternak, S., Liang, C., Zhang, J., Fulton, L., Graves, T.A., Minx, P., Reily, A.D., Courtney, L., Kruchowski, S.S., Tomlinson, C., Strong, C., Delehaunty, K., Fronick, C., Courtney, B., Rock, S.M., Belter, E., Du, F., Kim, K., Abbott, R.M., Cotton, M., Levy, A., Marchetto, P., Ochoa, K., Jackson, S.M., Gillam, B., Chen, W., Yan, L., Higginbotham, J., Cardenas, M., Waligorski, J., Applebaum, E., Phelps, L., Falcone, J., Kanchi, K., Thane, T., Scimone, A., Thane, N., Henke, J., Wang, T., Ruppert, J., Shah, N., Rotter, K., Hodges, J., Ingenthron, E., Cordes, M., Kohlberg, S., Sgro, J., Delgado, B., Mead, K., Chinwalla, A., Leonard, S., Crouse, K., Collura, K., Kudrna, D., Currie, J., He, R., Angelova, A., Rajasekar, S., Mueller, T., Lomeli, R., Scara, G., Ko, A., Delaney, K., Wissotski, M., Lopez, G., Campos, D., Braidotti, M., Ashley, E., Golser, W., Kim, H., Lee, S., Lin, J., Dujmic, Z., Kim, W., Talag, J., Zuccolo, A., Fan, C., Sebastian, A., Kramer, M., Spiegel, L., Nascimento, L., Zutavern, T., Miller, B., Ambroise, C., Muller, S., Spooner, W., Narechania, A., Ren, L., Wei, S., Kumari, S., Faga, B., Levy, M.J., McMahan, L., Van Buren, P., Vaughn, M.W., Ying, K., Yeh, C.-T., Emrich, S.J., Jia, Y., Kalyanaraman, A., Hsia, A.-P., Barbazuk, W.B., Baucom, R.S., Brutnell, T.P., Carpita, N.C., Chaparro, C., Chia, J.-M., Deragon, J.-M., Estill, J.C., Fu, Y., Jeddeloh, J.A., Han, Y., Lee, H., Li, P., Lisch, D.R., Liu, S., Liu, Z., Nagel, D.H., McCann, M.C., SanMiguel, P., Myers, A.M., Nettleton, D., Nguyen, J., Penning, B.W., Ponnala, L., Schneider, K.L., Schwartz, D.C., Sharma, A., Soderlund, C., Springer, N.M., Sun, Q., Wang, H., Waterman, M., Westerman, R., Wolfgruber, T.K., Yang, L., Yu, Y., Zhang, L., Zhou, S., Zhu, Q., Bennetzen, J.L., Dawe, R.K., Jiang, J., Jiang, N., Presting, G.G., Wessler, S.R., Aluru, S., Martienssen, R.A., Clifton, S.W., McCombie, W.R., Wing, R.A., Wilson, R.K., 2009. The B73 Maize Genome: Complexity, Diversity, and Dynamics. Science 326, 1112–1115. 10.1126/science.1178534

Serra Serra, N., Schanmuganathan, R., Becker, C. 2021. Allelopathy in rice: a story on momilactones, kin recognition, and weed management. J. Exp. Bot. erab084. 10.1093/jxb/erab084.

Shenton, M., Kobayashi, M., Terashima, S., Ohyanagi, H., Copetti, D., Hernández-Hernández, T., Zhang, J., Ohmido, N., Fujita, M., Toyoda, A., Ikawa, H., Fujiyama, A., Furuumi, H., Miyabayashi, T., Kubo, T., Kudrna, D., Wing, R., Yano, K., Nonomura, K.-I., Sato, Y., Kurata, N., 2020. Evolution and Diversity of the Wild Rice Oryza officinalis Complex, across Continents, Genome Types, and Ploidy Levels. Genome Biol. Evol. 12, 413–428. 10.1093/gbe/evaa037

Shi, C., Li, W., Zhang, Q.-J., Zhang, Y., Tong, Y., Li, K., Liu, Y.-L., Gao, L.-Z., 2020. The draft genome sequence of an upland wild rice species, Oryza granulata. Sci. Data 7, 131. 10.1038/s41597-020-0470-2

Shimura, K., Okada, A., Okada, K., Jikumaru, Y., Ko, K.-W., Toyomasu, T., Sassa, T., Hasegawa, M., Kodama, O., Shibuya, N., Koga, J., Nojiri, H., Yamane, H., 2007. Identification of a Biosynthetic Gene Cluster in Rice for Momilactones. J. Biol. Chem. 282, 34013–34018. 10.1074/jbc.M703344200

Smit, S.J., Lichman, B.R., 2022. Plant biosynthetic gene clusters in the context of metabolic evolution. Nat. Prod. Rep. 10.1039/D2NP00005A

Soreng, R.J., Peterson, P.M., Romaschenko, K., Davidse, G., Teisher, J.K., Clark, L.G., Barberá, P., Gillespie, L.J., Zuloaga, F.O., 2017. A worldwide phylogenetic classification of the Poaceae (Gramineae) II: An update and a comparison of two 2015 classifications. J. Syst. Evol. 55, 259–290. 10.1111/jse.12262

Sreedasyam, A., Plott, C., Hossain, M.S., Lovell, J.T., Grimwood, J., Jenkins, J.W., Daum, C., Barry, K., Carlson, J., Shu, S., Phillips, J., Amirebrahimi, M., Zane, M., Wang, M., Goodstein, D., Haas, F.B., Hiss, M., Perroud, P.-F., Jawdy, S.S., Yang, Y., Hu, R., Johnson, J., Kropat, J., Gallaher, S.D., Lipzen, A., Shakirov, E.V., Weng, X., Torres-Jerez, I., Weers, B., Conde, D., Pappas, M.R., Liu, L., Muchlinski, A., Jiang, H., Shyu, C., Huang, P., Sebastian, J., Laiben, C., Medlin, A., Carey, S., Carrell, A.A., Chen, J.-G., Perales, M., Swaminathan, K., Allona, I., Grattapaglia, D., Cooper, E.A., Tholl, D., Vogel, J.P., Weston, D.J., Yang, X., Brutnell, T.P., Kellogg, E.A., Baxter, I., Udvardi, M., Tang, Y., Mockler, T.C., Juenger, T.E., Mullet, J., Rensing, S.A., Tuskan, G.A., Merchant, S.S., Stacey, G., Schmutz, J., 2023. JGI Plant Gene Atlas: an updateable transcriptome resource to improve functional gene descriptions across the plant kingdom. Nucleic Acids Res. 51, 8383–8401. 10.1093/nar/gkad616

Stein, J.C., Yu, Y., Copetti, D., Zwickl, D.J., Zhang, L., Zhang, C., Chougule, K., Gao, D., Iwata, A., Goicoechea, J.L., Wei, S., Wang, J., Liao, Y., Wang, M., Jacquemin, J., Becker, C., Kudrna, D., Zhang, J., Londono, C.E.M., Song, X., Lee, S., Sanchez, P., Zuccolo, A., Ammiraju, J.S.S., Talag, J., Danowitz, A., Rivera, L.F., Gschwend, A.R., Noutsos, C., Wu, C., Kao, S., Zeng, J., Wei, F., Zhao, Q., Feng, Q., El Baidouri, M., Carpentier, M.-C., Lasserre, E., Cooke, R., Rosa Farias, D. da, da Maia, L.C., dos Santos, R.S., Nyberg, K.G., McNally, K.L., Mauleon, R., Alexandrov, N., Schmutz, J., Flowers, D., Fan, C., Weigel, D., Jena, K.K., Wicker, T., Chen, M., Han, B., Henry, R., Hsing, Y.C., Kurata, N., de Oliveira, A.C., Panaud, O., Jackson, S.A., Machado, C.A., Sanderson, M.J., Long, M., Ware, D., Wing, R.A., 2018. Genomes of 13 domesticated and wild rice relatives highlight genetic conservation, turnover and innovation across the genus Oryza. Nat. Genet. 50, 285–296. 10.1038/s41588-018-0040-0

Sun, G., Wase, N., Shu, S., Jenkins, J., Zhou, B., Torres-Rodríguez, J.V., Chen, C., Sandor, L., Plott, C., Yoshinga, Y., Daum, C., Qi, P., Barry, K., Lipzen, A., Berry, L., Pedersen, C., Gottilla, T., Foltz, A., Yu, H., O’Malley, R., Zhang, C., Devos, K.M., Sigmon, B., Yu, B., Obata, T., Schmutz, J., Schnable, J.C., 2022. Genome of Paspalum vaginatum and the role of trehalose mediated autophagy in increasing maize biomass. Nat. Commun. 13, 7731. 10.1038/s41467-022-35507-8

Tamogani, S., Mitani, M., Kodama, O., Akatsuka, T., 1993. Oryzalexin S structure: a new stemarane-type rice plant phytoalexin and its biogenesis. Tetrahedron 49, 2025–2032. 10.1016/S0040-4020(01)86302-X

Toyomasu, T., Goda, C., Sakai, A., Miyamoto, K., Shenton, M.R., Tomiyama, S., Mitsuhashi, W., Yamane, H., Kurata, N., Okada, K., 2018. Characterization of diterpene synthase genes in the wild rice species *Oryza brachyatha* provides evolutionary insight into rice phytoalexin biosynthesis. Biochem. Biophys. Res. Commun. 503, 1221–1227. 10.1016/j.bbrc.2018.07.028

Toyomasu, T., Miyamoto, K., Shenton, M.R., Sakai, A., Sugawara, C., Horie, K., Kawaide, H., Hasegawa, M., Chuba, M., Mitsuhashi, W., Yamane, H., Kurata, N., Okada, K., 2016. Characterization and evolutionary analysis of ent-kaurene synthase like genes from the wild rice species Oryza rufipogon. Biochem. Biophys. Res. Commun. 480, 402–408. 10.1016/j.bbrc.2016.10.062

Toyomasu, T., Shenton, M.R., Okada, K., 2020. Evolution of Labdane-Related Diterpene Synthases in Cereals. Plant Cell Physiol. 61, 1850–1859. 10.1093/pcp/pcaa106

Toyomasu, T., Tsukahara, M., Kaneko, A., Niida, R., Mitsuhashi, W., Dairi, T., Kato, N., Sassa, T., 2007. Fusicoccins are biosynthesized by an unusual chimera diterpene synthase in fungi. Proc. Natl. Acad. Sci. 104, 3084–3088. 10.1073/pnas.0608426104

VanBuren, R., Bryant, D., Edger, P.P., Tang, H., Burgess, D., Challabathula, D., Spittle, K., Hall, R., Gu, J., Lyons, E., Freeling, M., Bartels, D., Ten Hallers, B., Hastie, A., Michael, T.P., Mockler, T.C., 2015. Single-molecule sequencing of the desiccation-tolerant grass Oropetium thomaeum. Nature 527, 508–511. 10.1038/nature15714

Vogel, J.P., Garvin, D.F., Mockler, T.C., Schmutz, J., Rokhsar, D., Bevan, M.W., Barry, K., Lucas, S., Harmon-Smith, M., Lail, K., Tice, H., Schmutz (Leader), J., Grimwood, J., McKenzie, N., Bevan, M.W., Huo, N., Gu, Y.Q., Lazo, G.R., Anderson, O.D., Vogel (Leader), J.P., You, F.M., Luo, M.-C., Dvorak, J., Wright, J., Febrer, M., Bevan, M.W., Idziak, D., Hasterok, R., Garvin, D.F., Lindquist, E., Wang, M., Fox, S.E., Priest, H.D., Filichkin, S.A., Givan, S.A., Bryant, D.W., Chang, J.H., Mockler (Leader), T.C., Wu, H., Wu, W., Hsia, A.-P., Schnable, P.S., Kalyanaraman, A., Barbazuk, B., Michael, T.P., Hazen, S.P., Bragg, J.N., Laudencia-Chingcuanco, D., Vogel, J.P., Garvin, D.F., Weng, Y., McKenzie, N., Bevan, M.W., Haberer, G., Spannagl, M., Mayer (Leader), K., Rattei, T., Mitros, T., Rokhsar, D., Lee, S.-J., Rose, J.K.C., Mueller, L.A., York, T.L., Wicker (Leader), T., Buchmann, J.P., Tanskanen, J., Schulman (Leader), A.H., Gundlach, H., Wright, J., Bevan, M., Costa de Oliveira, A., da C. Maia, L., Belknap, W., Gu, Y.Q., Jiang, N., Lai, J., Zhu, L., Ma, J., Sun, C., Pritham, E., Salse (Leader), J., Murat, F., Abrouk, M., Haberer, G., Spannagl, M., Mayer, K., Bruggmann, R., Messing, J., You, F.M., Luo, M.-C., Dvorak, J., Fahlgren, N., Fox, S.E., Sullivan, C.M., Mockler, T.C., Carrington, J.C., Chapman, E.J., May, G.D., Zhai, J., Ganssmann, M., Guna Ranjan Gurazada, S., German, M., Meyers, B.C., Green (Leader), P.J., Bragg, J.N., Tyler, L., Wu, J., Gu, Y.Q., Lazo, G.R., Laudencia-Chingcuanco, D., Thomson, J., Vogel (Leader), J.P., Hazen, S.P., Chen, S., Scheller, H.V., Harholt, J., Ulvskov, P., Fox, S.E., Filichkin, S.A., Fahlgren, N., Kimbrel, J.A., Chang, J.H., Sullivan, C.M., Chapman, E.J., Carrington, J.C., Mockler, T.C., Bartley, L.E., Cao, P., Jung, K.-H., Sharma, M.K., Vega-Sanchez, M., Ronald, P., Dardick, C.D., De Bodt, S., Verelst, W., Inzé, D., Heese, M., Schnittger, A., Yang, X., Kalluri, U.C., Tuskan, G.A., Hua, Z., Vierstra, R.D., Garvin, D.F., Cui, Y., Ouyang, S., Sun, Q., Liu, Z., Yilmaz, A., Grotewold, E., Sibout, R., Hematy, K., Mouille, G., Höfte, H., Michael, T., Pelloux, J., O’Connor, D., Schnable, J., Rowe, S., Harmon, F., Cass, C.L., Sedbrook, J.C., Byrne, M.E., Walsh, S., Higgins, J., Bevan, M., Li, P., Brutnell, T., Unver, T., Budak, H., Belcram, H., Charles, M., Chalhoub, B., Baxter, I., The International Brachypodium Initiative, Principal investigators, DNA sequencing and assembly, Pseudomolecule assembly and BAC end sequencing, Transcriptome sequencing and analysis, Gene analysis and annotation, Repeats analysis, Comparative genomics, Small RNA analysis, Manual annotation and gene family analysis, 2010. Genome sequencing and analysis of the model grass Brachypodium distachyon. Nature 463, 763–768. 10.1038/nature08747

Wang, L., Zhu, T., Rodriguez, J.C., Deal, K.R., Dubcovsky, J., McGuire, P.E., Lux, T., Spannagl, M., Mayer, K.F.X., Baldrich, P., Meyers, B.C., Huo, N., Gu, Y.Q., Zhou, H., Devos, K.M., Bennetzen, J.L., Unver, T., Budak, H., Gulick, P.J., Galiba, G., Kalapos, B., Nelson, D.R., Li, P., You, F.M., Luo, M.-C., Dvorak, J., 2021. Aegilops tauschii genome assembly Aet v5.0 features greater sequence contiguity and improved annotation. G3 GenesGenomesGenetics 11, jkab325. 10.1093/g3journal/jkab325

Wickham, H., Averick, M., Bryan, J., Chang, W., McGowan, L.D., François, R., Grolemund, G., Hayes, A., Henry, L., Hester, J., Kuhn, M., Pedersen, T.L., Miller, E., Bache, S.M., Müller, K., Ooms, J., Robinson, D., Seidel, D.P., Spinu, V., Takahashi, K., Vaughan, D., Wilke, C., Woo, K., Yutani, H., 2019. Welcome to the Tidyverse. J. Open Source Softw. 4, 1686. 10.21105/joss.01686

Wilderman, P.R., Xu, M., Jin, Y., Coates, R.M., Peters, R.J., 2004. Identification of Syn-Pimara-7,15-Diene Synthase Reveals Functional Clustering of Terpene Synthases Involved in Rice Phytoalexin/Allelochemical Biosynthesis. Plant Physiol. 135, 2098–2105. 10.1104/pp.104.045971

Wu, D., Hu, Y., Akashi, S., Nojiri, H., Guo, L., Ye, C.-Y., Zhu, Q.-H., Okada, K., Fan, L., 2022a. Lateral transfers lead to the birth of momilactone biosynthetic gene clusters in grass. Plant J. 111, 1354– 1367. 10.1111/tpj.15893

Wu, D., Shen, E., Jiang, B., Feng, Y., Tang, W., Lao, S., Jia, L., Lin, H.-Y., Xie, L., Weng, X., Dong, C., Qian, Q., Lin, F., Xu, H., Lu, H., Cutti, L., Chen, H., Deng, S., Guo, L., Chuah, T.-S., Song, B.-K., Scarabel, L., Qiu, J., Zhu, Q.-H., Yu, Q., Timko, M.P., Yamaguchi, H., Merotto, A., Qiu, Y., Olsen, K.M., Fan, L., Ye, C.-Y., 2022b. Genomic insights into the evolution of Echinochloa species as weed and orphan crop. Nat. Commun. 13, 689. 10.1038/s41467-022-28359-9

Xu, M., Hillwig, M.L., Prisic, S., Coates, R.M., Peters, R.J., 2004. Functional identification of rice syn-copalyl diphosphate synthase and its role in initiating biosynthesis of diterpenoid phytoalexin/allelopathic natural products. Plant J. 39, 309–318. 10.1111/j.1365-313X.2004.02137.x

Yan, N., Yang, T., Yu, X.-T., Shang, L.-G., Guo, D.-P., Zhang, Y., Meng, L., Qi, Q.-Q., Li, Y.-L., Du, Y.-M., Liu, X.-M., Yuan, X.-L., Qin, P., Qiu, J., Qian, Q., Zhang, Z.-F., 2022. Chromosome-level genome assembly of Zizania latifolia provides insights into its seed shattering and phytocassane biosynthesis. Commun. Biol. 5, 1–11. 10.1038/s42003-021-02993-3

Yang, X., Gao, S., Guo, L., Wang, B., Jia, Y., Zhou, J., Che, Y., Jia, P., Lin, J., Xu, T., Sun, J., Ye, K., 2021. Three chromosome-scale Papaver genomes reveal punctuated patchwork evolution of the morphinan and noscapine biosynthesis pathway. Nat. Commun. 12, 6030. 10.1038/s41467-021-26330-8

Ye, Z., Nakagawa, K., Natsume, M., Nojiri, H., Kawaide, H., Okada, K. 2017. Biochemical synthesis of uniformly ^13^C-labeled diterpene hydrocarbons and their bioconversion to diterpenoid phytoalexins in planta. Biosci. Biotechnol. Biochem. 81, 1176–1184. 10.1080/09168451.2017.1285689

Ye, C.-Y., Wu, D., Mao, L., Jia, L., Qiu, J., Lao, S., Chen, M., Jiang, B., Tang, W., Peng, Q., Pan, L., Wang, L., Feng, X., Guo, L., Zhang, C., Kellogg, E.A., Olsen, K.M., Bai, L., Fan, L., 2020. The Genomes of the Allohexaploid Echinochloa crus-galli and Its Progenitors Provide Insights into Polyploidization-Driven Adaptation. Mol. Plant 13, 1298–1310. 10.1016/j.molp.2020.07.001

Yu, H., Lin, T., Meng, X., Du, H., Zhang, J., Liu, G., Chen, Mingjiang, Jing, Y., Kou, L., Li, X., Gao, Q., Liang, Yan, Liu, X., Fan, Z., Liang, Yuntao, Cheng, Z., Chen, Mingsheng, Tian, Z., Wang, Y., Chu, C., Zuo, J., Wan, J., Qian, Q., Han, B., Zuccolo, A., Wing, R.A., Gao, C., Liang, C., Li, J., 2021. A route to de novo domestication of wild allotetraploid rice. Cell 184, 1156–1170.e14. 10.1016/j.cell.2021.01.013

Zhang, J., Wu, F., Yan, Q., John, U.P., Cao, M., Xu, P., Zhang, Z., Ma, T., Zong, X., Li, J., Liu, R., Zhang, Y., Zhao, Y., Kanzana, G., Lv, Y., Nan, Z., Spangenberg, G., Wang, Y., 2021. The genome of Cleistogenes songorica provides a blueprint for functional dissection of dimorphic flower differentiation and drought adaptability. Plant Biotechnol. J. 19, 532–547. 10.1111/pbi.13483

Zhao, H., Gao, Z., Wang, L., Wang, J., Wang, Songbo, Fei, B., Chen, C., Shi, C., Liu, X., Zhang, H., Lou, Y., Chen, L., Sun, H., Zhou, X., Wang, Sining, Zhang, C., Xu, H., Li, L., Yang, Y., Wei, Y., Yang, W., Gao, Q., Yang, H., Zhao, S., Jiang, Z., 2018. Chromosome-level reference genome and alternative splicing atlas of moso bamboo (Phyllostachys edulis). GigaScience 7, giy115. 10.1093/gigascience/giy115

Zhao, H., Wang, W., Yang, Y., Wang, Z., Sun, J., Yuan, K., Rabbi, S.M.H.A., Khanam, M., Kabir, M.S., Seraj, Z.I., Rahman, M.S., Zhang, Z., 2023. A high-quality chromosome-level wild rice genome of Oryza coarctata. Sci. Data 10, 701. 10.1038/s41597-023-02594-1

Zhou, Y., Ma, Y., Zeng, J., Duan, L., Xue, X., Wang, H., Lin, T., Liu, Z., Zeng, K., Zhong, Y., Zhang, S., Hu, Q., Liu, M., Zhang, H., Reed, J., Moses, T., Liu, Xinyan, Huang, P., Qing, Z., Liu, Xiubin, Tu, P., Kuang, H., Zhang, Z., Osbourn, A., Ro, D.-K., Shang, Y., Huang, S., 2016. Convergence and divergence of bitterness biosynthesis and regulation in Cucurbitaceae. Nat. Plants 2, 1–8. 10.1038/nplants.2016.183

Zi, J., Mafu, S., Peters, R.J., 2014. To Gibberellins and Beyond! Surveying the Evolution of (Di)Terpenoid Metabolism. Annu. Rev. Plant Biol. 65, 259–286. 10.1146/annurev-arplant-050213-035705

